# Knockout of endoplasmic reticulum localized molecular chaperone HSP90.7 impairs seeding development and cellular auxin homeostasis in Arabidopsis

**DOI:** 10.1101/2023.03.06.531358

**Authors:** Jenan Noureddine, Bona Mu, Homaira Hamidzada, Wai Lam Mok, Diana Bonea, Eiji Nambara, Rongmin Zhao

## Abstract

The Arabidopsis endoplasmic reticulum localized heat shock protein HSP90.7 modulates tissue differentiation and stress responses; however, complete knockout lines have not been previously reported. In this study, we identified and analyzed a mutant allele, *hsp90.7-1*, which did not express any protein and showed seedling lethality. Microscopic analyses revealed its essential role in male and female fertility, trichomes and root hairs development, proper chloroplast function, and in apical meristem maintenance and differentiation. Comparative transcriptome and proteome analyses also revealed a role of the protein in a multitude of cellular processes. Particularly, the auxin responsive pathway was specifically down-regulated in the *hsp90.7-1* mutant seedlings. We measured a much-reduced auxin content in both root and shoot tissues. Through comprehensive histological and molecular analyses, we demonstrated PIN1 and PIN5 expressions were dramatically reduced in the mutant, and the TAA-YUCCA primary auxin biosynthesis pathway was also down-regulated, thus revealing a critical new role of HSP90.7 in the regulation of auxin responses. This study therefore not only fulfilled a gap in understanding the essential role of HSP90 paralogs in eukaryotes, but also provided a mechanistic insight on this molecular chaperone in regulating plant growth and development via modulating cellular auxin homeostasis.

## INTRODUCTION

Arabidopsis HSP90.7 is an endoplasmic reticulum (ER) lumen localized HSP90 family heat shock protein in plants, and also termed SHEPHERD, as previous studies on a T-DNA insertion line revealed its potential role in regulating the function of CLAVATA proteins though the exact target is unknown (Ishiguro *et al*., 2002). In higher organisms, HSP90 proteins are divided into five subfamilies based on their structure and subcellular location (Chen *et al*., 2006). Cytosolic HSP90A is absolutely required in eukaryotes, and its co-chaperones and many client proteins have been well studied (Prodromou, 2016, Biebl and Buchner, 2019). Organellar HSP90 homologues generally lack co-chaperones in comparison with the cytosolic form, and instead, their intrinsically extra but unique structure helps to accomplish the functional cycle as exemplified in mammalian HSP90B/GRP94 and TRAP1 (Elnatan *et al*., 2017, Huck *et al*., 2017). Some co-chaperones may assist GRP94 function, including MZB1 which may assist GRP94 in folding µ heavy chain (µHC) during ER stress (Rosenbaum *et al*., 2014), CNPY3 in folding TLRs (Liu *et al*., 2010), and the closure accelerating cochaperone BiP, an HSP70 family protein (Huang *et al*., 2022) which binds to GRP94 N-terminal and middle domain (Sun *et al*., 2019). Bacterial HtpG (Wong and Houry, 2004) and mammalian mitochondrial TRAP1 (Lisanti *et al*., 2014) are not essential for the cell viability or animal development under normal condition. HSP90B/GRP94 in mammals and HSP90C in plants, however, are essential and knockout lines show embryonic lethality (Mao *et al*., 2010, Inoue *et al*., 2013). Nevertheless, whether the plant ER-localized GRP94, e.g. HSP90.7 in Arabidopsis, is absolutely essential or not is still unclear as a complete knockout line is not yet reported.

HSP90.7 is particularly important for proliferation tissues to support protein secretion (Klein *et al*., 2006), and it has a short charged region in the middle domain that plays a critical role in regulating ATP hydrolysis and ER stress resistance (Chong *et al*., 2015). Overexpression of ER-localized HSP90.7 alleviated the ER stress under high concentrations of calcium (Song *et al*., 2009). It should be noted, however, the *shepherd* mutant was originally identified from the Wassilewskija ecotype, analyzed by crossing with *clavata* and *wuschel* mutants in the Landsberg *erecta* ecotype (Ishiguro *et al*., 2002), and still appeared leaky expression (Klein *et al*., 2006). Additionally, while some candidate HSP90.7 interactors have been previously identified by yeast two-hybrid assays (Chong *et al*., 2015), direct evidence for HSP90.7 to support a native client protein folding or targeting in vivo is still lacking.

Auxin is a key plant growth hormone with a major role in stimulating growth and it regulates almost every aspect of cellular processes (Leyser, 2018). For instance, auxin forms a gradient in shoot apical meristem (Vernoux *et al*., 2010) and regulates organ differentiation together with the WUSCHEL-CLAVATA pathway (Somssich *et al*., 2016). Indole-3-acetic acid (IAA) is the predominant form of naturally occurring auxin and it acts as a molecular glue to stabilize the interaction between the SCF E3 ligase subunit F-box protein TIR1 and the transcription repressor substrates Aux/IAA (Tan *et al*., 2007), thus promoting the degradation of those transcription repressors. Consequently, the auxin response factors (ARFs) are released from the ARF/IAA complex to induce the expression of a large set of auxin responsive genes (Hagen and Guilfoyle, 2002). Auxin is not homogenously distributed within cells and tissues and its regulatory role is controlled by a complex auxin homeostasis system (Ljung, 2013) involving auxin biosynthesis (Kasahara, 2016), catabolism (Stepanova and Alonso, 2016), long and short distance transport via a group of auxin transporters including PIN proteins, which are responsible for polar auxin transport (PAT) (Krecek *et al*., 2009). However, how proteins regulating auxin homeostasis themselves are regulated at the post-translational levels are generally less studied. It has been reported that cytosolic HSP90 stabilizes TIR1 protein in Arabidopsis (Wang *et al*., 2016). HSP90 co-chaperone p23 regulates root auxin distribution (D’Alessandro *et al*., 2015). PIN proteins on plasma membrane can be internalized by endocytosis and recycled upon treatment with trafficking inhibitor (Kleine-Vehn and Friml, 2008), natural or synthetic auxins (Narasimhan *et al*., 2021), signifying the complex regulation network of auxin response. Nevertheless, how molecular chaperones within the secretory pathway regulate the folding and targeting of these critical PIN proteins are still largely unknown.

In this study, we identified a new Arabidopsis T-DNA insertion knockout line *hsp90.7-1* in Columbia ecotype, in which HSP90.7 protein is not detectable. The development of *hsp90.7-1* homozygous mutant was arrested at the seedling stage. Comprehensive microscopic analyses revealed significantly impaired organization of the shoot and root meristems. We also performed RNA sequencing and proteomics analyses to identify global gene expression and protein profile differences in the mutant seedlings. It turned out that the auxin responsive pathway was one of the only few that were downregulated in the mutant. We measured a much-reduced auxin content in both root and shoot tissues of the mutant. We also showed that PIN1 and PIN5 expressions were dramatically reduced, and the TAA-YUCCA primary auxin biosynthesis pathway was down-regulated as well in the mutant. Our study therefore not only provided evidence showing the global role of the molecular chaperone HSP90.7, but also provided a mechanistic insight on the ER-localized chaperone in regulating plant growth and development via modulating cellular auxin homeostasis.

## RESULTS

### Knockout of ER-localized HSP90.7 caused an early developmental arrest in Arabidopsis

To understand the physiological function of HSP90.7, we requested a few T-DNA insertional SALK and SAIL lines from ABRC and confirmed a T-DNA insertion allele in CS825167, which we refer to as *hsp90.7-1* in this study. The T-DNA was inserted in the second exon, disrupting translation of the protein from Ala-28 (Figure 1a, Figure S1a), and the Basta resistance gene in T-DNA was closely associated with the mutation after four times backcrossing (Figure S1b and 1c). Analysis of self-pollinated and backcrossed progenies indicated that *hsp90.7-1* allele did not show Mendelian segregation and only approximately 10% *hsp90.7-1* pollens reached the ovules for successful pollination, suggesting partially male sterile and very rarely observed mutant homozygous seedlings (Figure 1b, Table 1, Table S1). These resemble the *shepherd* mutant showing poor male fertility (Ishiguro *et al*., 2002). However, different from *shepherd* that still has a low level HSP90.7 expression, grows to flowering stage and is partially fertile, *hsp90.7-1* homozygous mutant is arrested at the early seedling stage and lacks proper true leaf development (Figure 1b). Additionally, mutant cotyledons are over-bent and appear dark green to purple in color, and the HSP90.7 protein is not detectable at all (Figure 1b).

**Figure 1.**
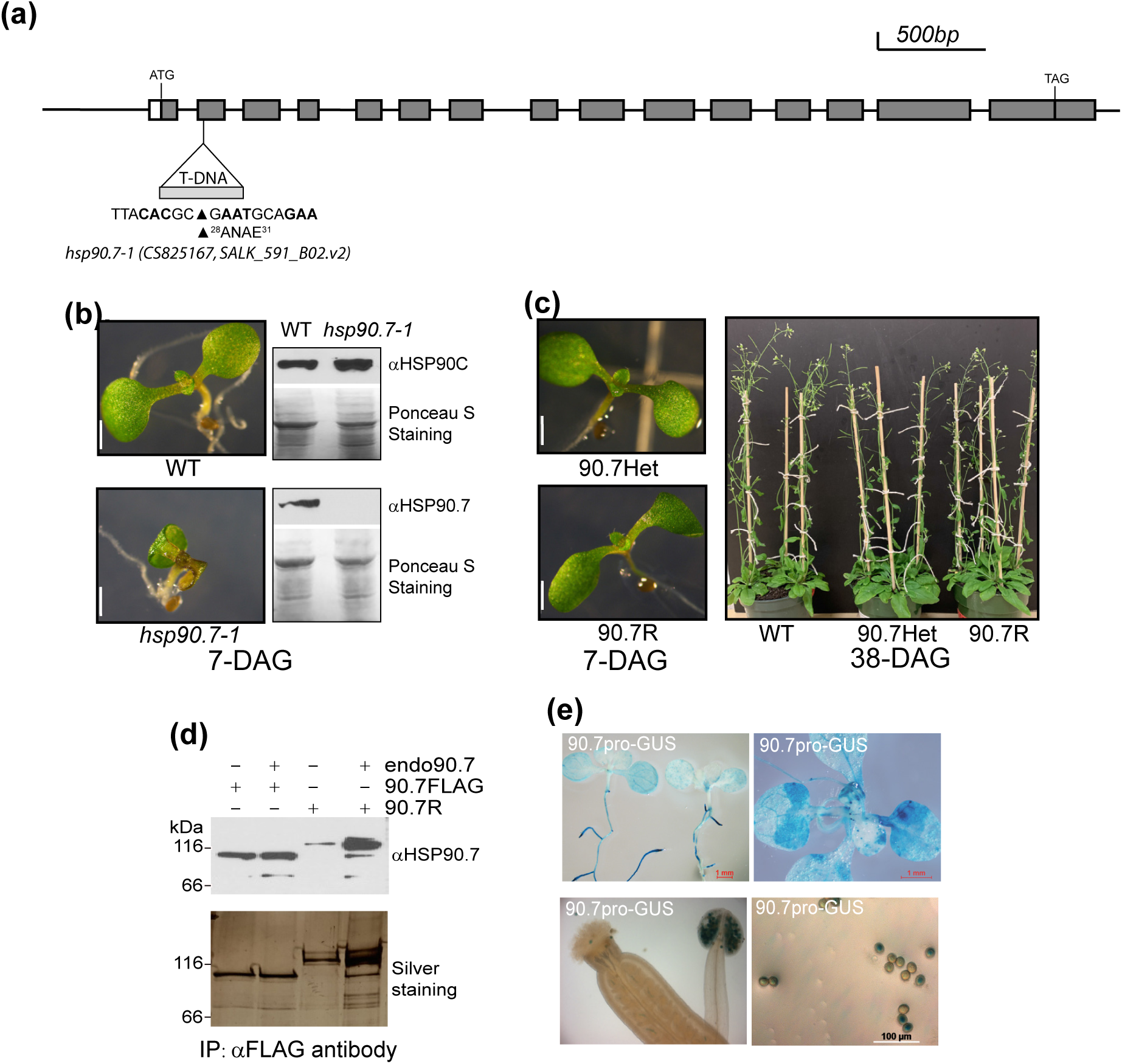
HSP90.7 knockout mutant exhibits significant growth defects. (a) Schematic diagram of T-DNA insertion mutant alleles in the *HSP90.7* gene. *hsp90.7-1* allele carries a T-DNA inserted inside the second exon as well as a Basta resistant gene. (b) Seedlings grown at 7 days after germination (DAG). On the right are the immunoblots of total cellular proteins with anti-HSP90.7 and anti-HSP90C antibodies. (c) Plants grown at 7-DAG and 38-DAG. Note the labeling of 90.7Het for *hsp90.7-1* heterozygote, 90.7F for FLAG-tagged HSP90.7 protein expressed from its native promoter, and 90.7R for a mCherry- and FLAG-tagged HSP90.7 protein expressed from its native promoter are used throughout all figures unless specifically indicated. (d) Immunoblots (top) and silver staining (bottom) of anti-FLAG antibody affinity gel purified proteins from different transgenic plant lines. (e) GUS staining of seedlings and pollens from HSP90.7pro-GUS transgenic plants.

**Table 1:** Segregation ratio of HSP90.7 mutants.

| **Tab** | **Progeny** |  |  |  |
| --- | --- | --- | --- | --- |
| **Crosses** | *HSP90.7*<sup>-/-</sup> (*aa*) | *HSP90.7*<sup>+/-</sup> (*Aa*) | *HSP90.7*<sup>+/+</sup> (*AA*) | **Total # of Seeds** |
| WT self | 0 (0) | 0 (0) | 594 (549) | 594 |
| *HSP90.7*<sup>+/-</sup> self * | 31 (160.25) | 287 (310.5) | 325 (160.25) | 641 |
| WT ♀ X *HSP90.7*<sup>+/-</sup> ♂ * | 0 (0) | 17 (182.5) | 348 (182.5) | 365 |
| *HSP90.7*<sup>+/-</sup> ♀ X WT ♂ | 0 (0) | 235 (236.5) | 238 (236.5) | 473 |

Note: Numbers in bracket represent expected numbers. \* Represents the observed number is significantly different from expected number with $p < 0.05$ by $\chi^2$ -test.

To further confirm that the arrested growth phenotype only resulted from *hsp90.7-1* mutation, we generated a FLAG-tagged HSP90.7 construct driven by the endogenous HSP90.7 promoter, and a similar construct in which HSP90.7-FLAG is fused to an mCherry tag. Both HSP90.7 fusion constructs complemented the growth defects of the *hsp90.7-1* plants (Figure 1c,d) and rescued the poor male fertility phenotype (Table S1). This also indicates fusion of a FLAG or a larger mCherry tag does not interfere with the *in vivo* function of HSP90.7 for vegetative shoot growth. We also attempted to rescue the *hsp90.7-1* mutant phenotype by re-introducing the FLAG-tagged HSP90.7 protein expressed under the CaMV 35S promoter (Chong *et al*., 2015). It was surprisingly noted that HSP90.7 driven by the CaMV 35S promoter did not rescue the poor male fertility phenotype (Table S1). We hypothesized that this may be due to the poor activity of CaMV 35S promoter in Arabidopsis pollens as previously reported (Wilkinson *et al*., 1997) while HSP90.7 is highly active in pollens. To confirm this, we made an HSP90.7pro:GUS reporter line. The expression of *HSP90.7* was well observed in seedling root and shoot tissues as well as pollen grains (Figure 1e). We also investigated the expression profile of HSP90.7 using microarray data and showed that HSP90.7 is widely expressed (Figure S2), generally in agreement with the GUS reporter analysis except GUS reporter analyses indicated a high expression of HSP90.7 in pollens.

Interestingly, heterozygote flowers appeared to have very similar overall development to wild type, except more aberrant smaller and shrunken pollens (Figure 2a). The heterozygotes also had much lower seed setting and many unfertilized ovules (Figure 2b,c) although the total number of ovules within the pistils were not much different (Figure 2d). The siliques on the primary inflorescence were significantly shorter than wild type (Figure 2e), a typical phenotype associated with defects in pollen tube growth and/or female gametophyte development (Madison *et al*., 2015, Sato and Maeshima, 2019). We then examined pollen germination in vitro and indeed observed many pollens from heterozygote plants, likely emerging from mutant pollens, did not germinate well (Figure 2f,g). We hypothesize that mutant pollen tubes within the pistil may elongate slowly and primarily reach ovules closer to the stigma. To test this, we examined seeds collected separately from the top and the bottom sections of a mature silique, and as expected, we observed an overall more Basta resistant seeds corresponding to the T-DNA insertion mutants in the top region (Figure 2h) as well as more occurrence of mutant homozygotes (Table 2).

**Figure 2.**
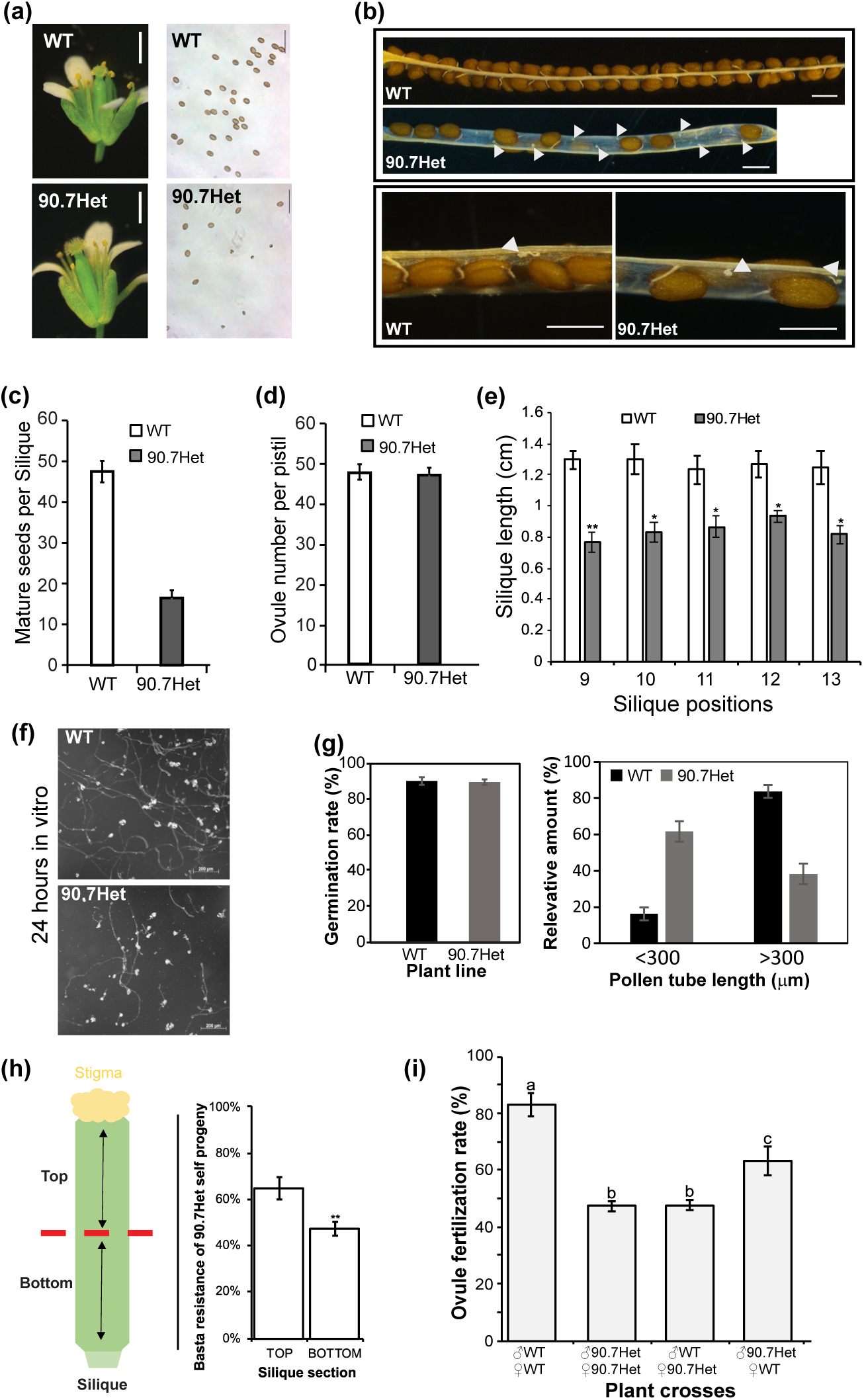
Fertility of *hsp90.7-1* heterozygote plants. (a) wild type (WT) and 90.7Het flowers (left) and mature pollens (right). (b) Mature siliques from wildtype and 90.7Het plants. Arrows indicate unfertilized ovules. (c-e) Average mature seed number, total ovules and silique lengths of wildtype and 90.7Het plants. (f) Images of pollen tubes from WT and 90.7Het pollens germinated in vitro for 24 hours. (g) Pollen germination rate after 24 hours in vitro germination (left), and the pollen tube length distribution (right). (h) Percentage of Basta resistant seeds from top and bottom halves of pistils from self-pollinated 90.7Het plants. (i) Ovule fertilization rates from different crosses with 90.7Het plants. Different letters on the top of the error bars represent difference between the two groups are statistically significant.

**Table 2:**
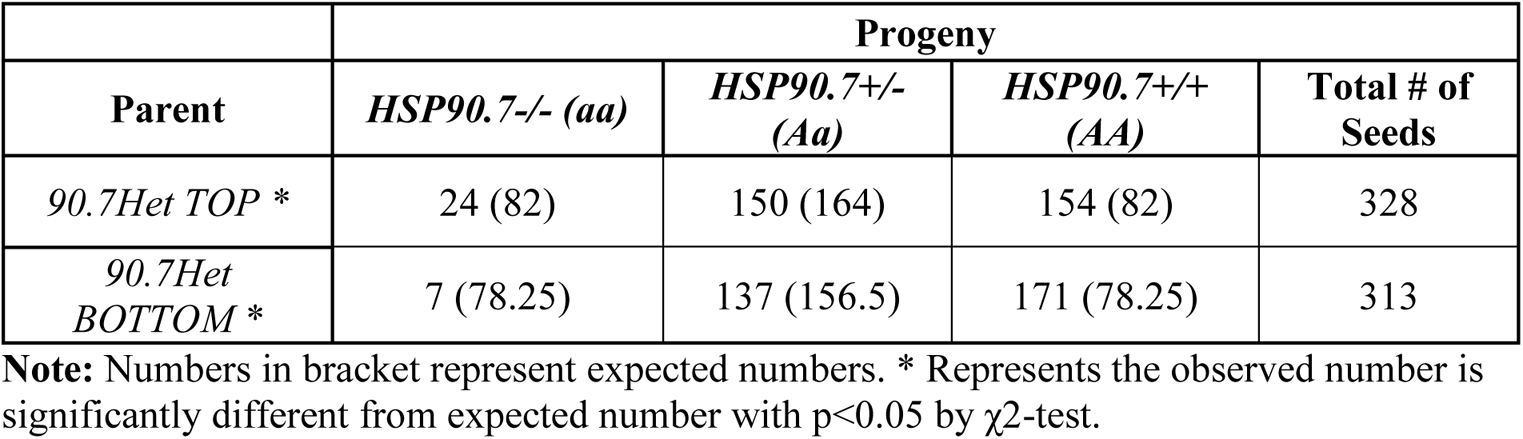
Top and bottom section of heterozygote siliques.

|  | **Progeny** |  |  |  |
| --- | --- | --- | --- | --- |
| **Parent** | *HSP90.7*<sup>-/-</sup> (*aa*) | *HSP90.7*<sup>+/-</sup> (*Aa*) | *HSP90.7*<sup>+/+</sup> (*AA*) | **Total # of Seeds** |
| *90.7Het TOP* * | 24 (82) | 150 (164) | 154 (82) | 328 |
| *90.7Het BOTTOM* * | 7 (78.25) | 137 (156.5) | 171 (78.25) | 313 |

**Note:** Numbers in bracket represent expected numbers. \* Represents the observed number is significantly different from expected number with $p < 0.05$ by $\chi^2$ -test.

Plant fertility could be affected by a variety of factors, and defective female gametophyte development (Shimizu and Okada, 2000) and/or defective transmission duct tissues (Wu *et al*., 1995) also impair the pollen tube funiculus and micropyle guidance. To understand whether the low fertility rate was solely caused by the slow pollen tube elongation, we examined the F1 siliques of different crosses, and noticed that, when the paternal plant was wild type, the overall seed setting of the heterozygote resembled that of self-pollinated heterozygote plants (Figure 2i). This suggests *hsp90.7-1* mutant also had defective female fertility.

### HSP90.7 is essential for root meristem cell proliferation and expansion

We examined *hsp90.7-1* mutants for their overall root growth. It is evident that the mutant had a significant reduction in primary root growth and elongation and re-introducing mCherry-tagged HSP90.7 completely rescued the growth defect (Figure S3a,b). We also observed a significant increase in root hair density in the mutants (Figure S3c). The mutant central cells and surrounding initials were difficult to identify compared to the wild type (Figure S3d). There were significantly less cortical cells that were also smaller in size in the meristem zone for the mutant roots as counted along the longitudinal plane, and not surprisingly, mCherry tagged HSP90.7 rescued the mutant to wild type phenotype (Figure S3f-g).

Next, we took advantage of a cohort of well-characterized cell division and root meristem specific maker lines including *QC25GUS*, a quiescent center promoter trap line (Sabatini *et al*., 1999) and *CYCB1-1GUS,* a cell division maker which accumulates in cells with mitotic potential and involved in growth control at the G2/M phase (Colon-Carmona *et al*., 1999). Interestingly, both *QC25GUS* and *CYCB1-1GUS* expression appeared to be significantly repressed in the mutant (Figure 3a), showing the quiescent center was severely impaired and cells prematurely exiting the cell cycle in the root tip region. We also analyzed the expression for *SHRGFP* and *J2341GFP* which are all uniquely expressed in specific root meristem cells or tissues (Garcia-Gomez *et al*., 2021), but no change of their tissue-specific or subcellular localization, except a much lower expression of SHRGFP signal (Figure 3a), suggesting cell patterning was greatly impaired in the *hsp90.7-1* root tip. WOX5 localizes at the QC and represses cell division to maintain its quiescence (Pi *et al*., 2015). Surprisingly, *WOX5GFP* in *hsp90.7-1* was not much changed (Figure 3a).

**Figure 3.**
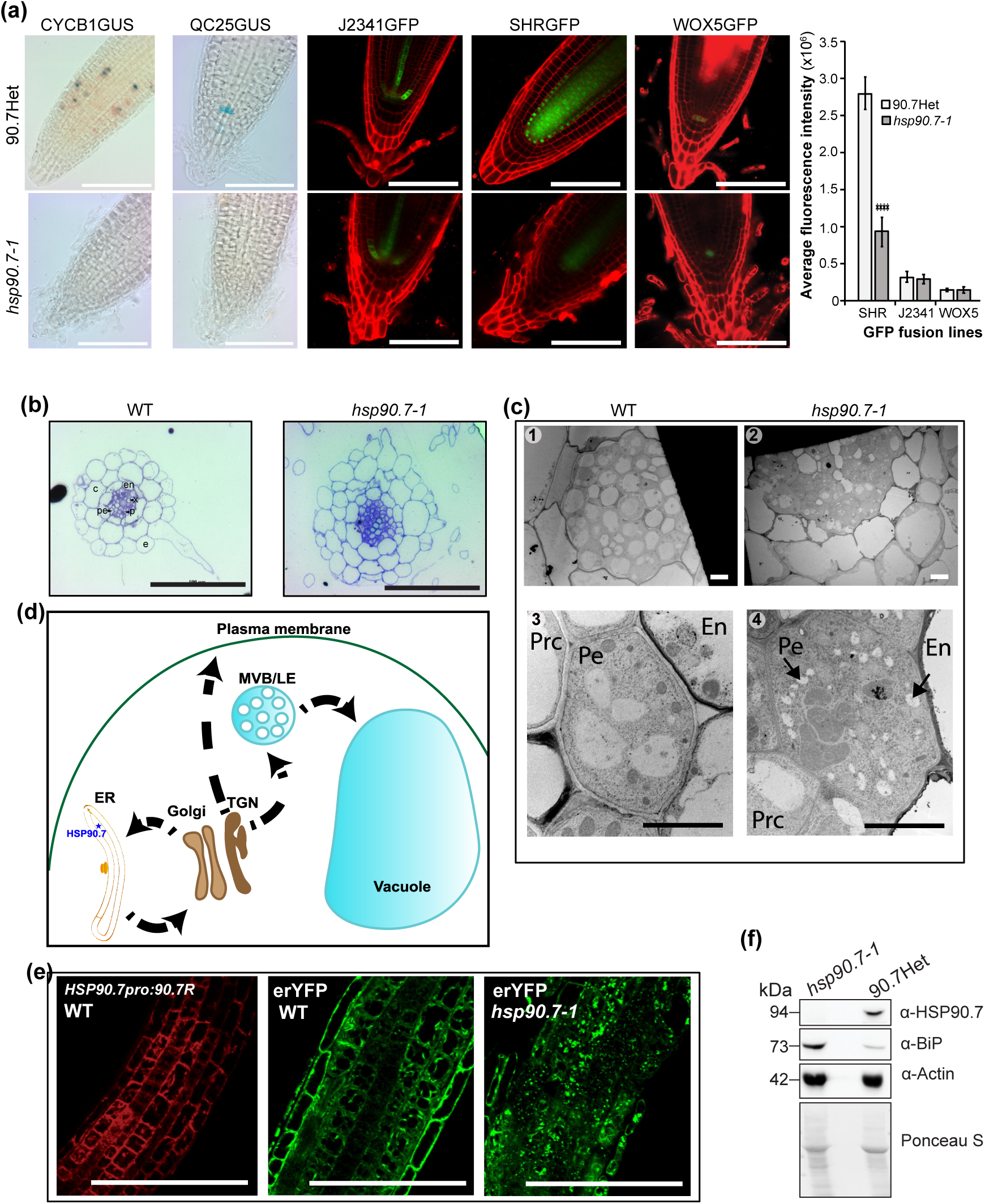
HSP90.7 is essential for root development. (a) Left, GUS staining and confocal microscopic GFP fluorescent images of root tip tissues expressing CYCB1;1GUS, QC25GUS, J2341GFP, SHRGFP or WOX5GFP marker proteins. Right, quantitative analysis of GFP fluorescent signals using Image J. root tissues were also stained with propidium iodide to show the border of cells. Scale bar, 100 μm. Error bars represent standard deviation and Student’s t-test was used for statistical analysis. **** indicates P < 0.00005. (b) Histological analysis of primary root maturation zone of wild type and *hsp90.7-1* mutant roots. Scale bar, 100 μm. e, epidermis; c, cortex; en, endodermis; pe, pericycle; x, xylem; p, phloem. (c)Transmission electron microscopic images of 6-DAG wild type and *hsp90.7-1* mutant root maturation zone cells. Pe, pericycle cells; En, endodermal cells; Prc, procambium. Scale bar, 5 μm. (d) Schematic diagram of the plant secretion pathway. Co-translational translocation of secreted proteins into the endoplasmic reticulum (ER) is followed by transport via ER-derived vesicles to the Golgi apparatus and the trans-Golgi network (TGN). From the TGN, cargo is transported to the plasma membrane or to the multivesicular bodies (MVBs)/late endosomes (LEs) and towards the vacuole. HSP90.7 is indicated by a blue star. [Adapted from (Ruano and Scheuring, 2020)] (e) Confocal images of mCherry fluorescence from 90.7R transgenic root elongation zone cells (left), and ER-located YFP fluorescence in wild type and *hsp90.7-1* root elongation zone tissues (middle and right). Scale bar, 100 μm. (f) Immunoblot with anti-HSP90.7, anti-BiP and anti-Actin antibodies for 90.7Het and *hsp90.7-1* 8-DAG seedlings total proteins.

In Arabidopsis, the wildtype root exhibits radial symmetry and contains an invariable number of 8 cortical and endodermal cells (Figure 3b). It is evident that the *hsp90.7-1* mutant roots lost the radial symmetry and had more cortical and endodermal cells in the maturation zone (Figure 3b). This suggests abnormal cell division and patterning in the mutant. Additionally, mutant cells were generally swelled and enlarged, and the layers of epidermis, cortex and endodermis were not clearly separated. We also observed the ultrastructure of *hsp90.7-1* tissues using transmission electron microscopy. Many small low electron density oval shaped structures appeared in the *hsp90.7-1* mutant vasculature cells (Figure 3c, arrow heads). It is likely that these smaller oval structures were premature vacuoles or some other ER-related or ER-originated vesicles (Figure 3d). We introduced an ER-lumen located YFP marker to the mutant and observed that the ER-localized YFP protein was distributed very differently and in many more vesicular structures, while the distribution of the ER-located YFP and HSP90.7-mCherry were comparable in wildtype plant (Figure 3e), suggesting abnormal cellular ER network distribution and/or abnormal accumulation of ER-originated vesicles. BiP is an ER lumen located HSP70 family heat shock protein, and its up regulation is commonly used as a hallmark for ER stress (Gardner *et al*., 2013). Similarly, immunoblot analysis indicated the *hsp90.7-1* mutant had accumulated more BiP protein (Figure 3f).

### HSP90.7 is essential for shoot meristem cell proliferation and expansion

To have a close look at the apical meristem tissues, we sectioned the apical meristem and observed that the shoot apical meristem (SAM) in *hsp90.7-1* seedlings appeared more dome shaped compared to the wild type (Figure 4a). The mutant SAM cells also appear larger in size, a wider diameter compared to the wild type, an indication of disturbed meristem homeostasis and delayed leaf primordia differentiation (Mandel *et al*., 2016). Scanning electron microscopy showed that the true leaf development was significantly affected, with abnormal trichome initiation and maturation (Figure 4b). We then analyzed the expression of a few genes in the WUSCHEL-CLAVATA pathway primarily controlling the apical meristem niche. It was shown that the expression of differentiation promoting *CLV3* was increased and the kinase receptor *CLV1* gene was decreased, while the expression of another short *CLV2* gene and the transcription factor *WUS* gene were not significantly affected (Figure 4c). This indicates CLV1 might be the major protein affected in the mutant. These microscopic and gene expression analyses clearly indicate that HSP90.7 is required for differentiation of shoot tissues, thus elucidating a mechanism of seedling lethality phenotype. It should be noted that many mutant shoot cells accumulated abnormal starch granules in chloroplasts (Figure 4a), and the abnormal accumulation and breakdown of starch granules were also confirmed by transmission electron microscopy analysis of the cotyledon cells (Figure 4d). This may explain the dark green color of the cotyledons from *hsp90.7-1* mutant seedlings (Figure 1b). This also indicates HSP90.7 may play a global role in many other cellular processes.

**Figure 4.**
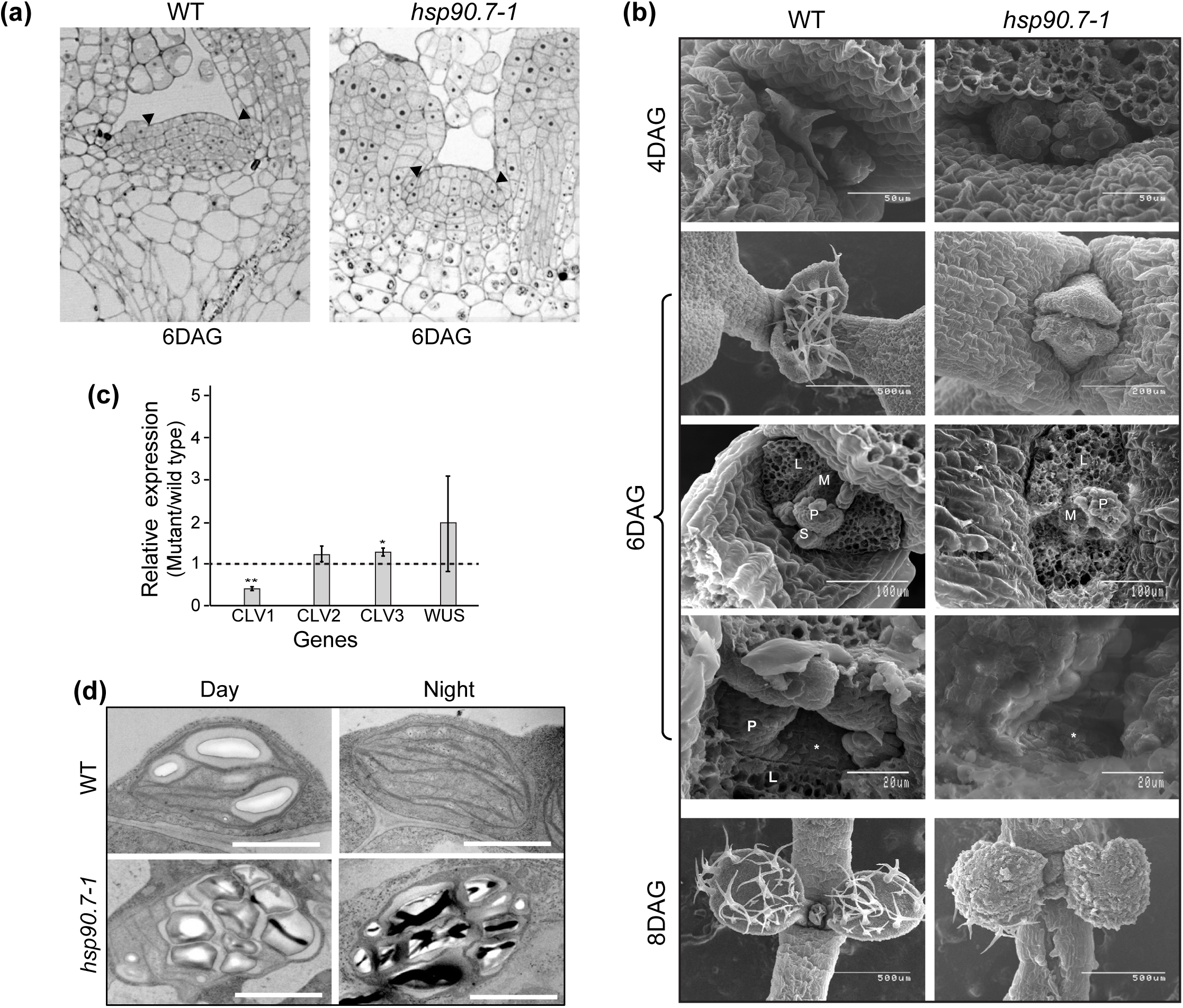
HSP90.7 knockout mutant exhibits impaired shoot differentiation and abnormal starch granule accumulation. (a) Longitudinal sections of the shoot apical meristem (SAM) from 6-DAG seedlings. Black arrow heads indicate borders of the peripheral zone. Scale bar: 100 µm. (b) Scanning electron microscopic images of shoot apex from 4, 6 and 8-DAG seedlings. M, shoot apex; P, leaf primordia; S, stipules; L, leaf trace. (c) RT-qPCR analysis of *CLV1*, *CLV2*, *CLV3* and *WUS* gene expression in WT and *hsp90.7-1* 8-DAG seedlings. ** indicates P < 0.005. (d) Transmission electron microscopic images of cotyledon palisade cells from 6-DAG seedlings. Day and Night represent samples collected at the end and beginning of the photoperiod, respectively. Scale bar: 5 µm.

### Knockout of HSP90.7 affects global gene expression and protein enrichment profiles

To gain insight on global gene expression patterns in the *hsp90.7-1* mutant relative to wild type, we conducted a comparative transcriptome analysis using bulk RNA sequencing of 8-day-old seedlings. A total of 17600 genes were detected, with 1516 genes differentially up-regulated, and 900 genes down-regulated by at least 2 folds (Figure 5a, Table S2). It is interesting to note that HSP90.7 gene itself was not recognized as a significant differentially expressed gene at the transcriptional level, however, the reading frame of *HSP90.7* gene was clearly disrupted on the second exon (Figure S4), supporting our previous immunoblot analysis (Figure 1b).

**Figure 5.**
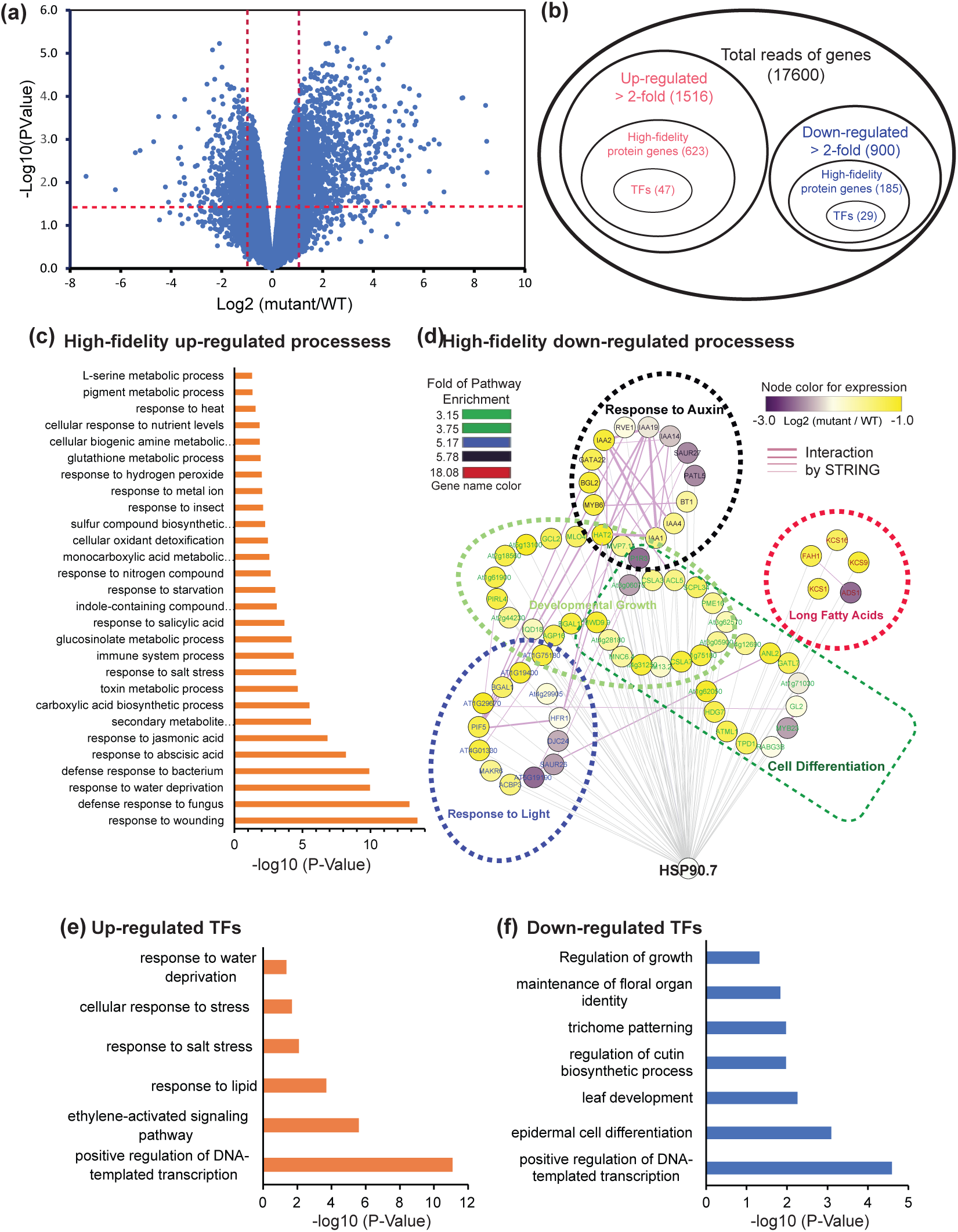
Summary of comparative transcriptome analyses of wild type and mutant *hsp90.7-1* seedlings. (a) Volcano plot showing DEGs between the mutant and wild type seedlings as revealed by RNA sequencing. (b) Summary of total identified genes and the DEGs. The raw data are indicated in Table S2. High-fidelity protein genes represent those that were altered at least 2-fold with P-value <0.05 and false discovery rate <0.05. (c) Significantly up-regulated processes based on high fidelity core set of DEGs (b). GO terms were generated with PANTHER overrepresentation test by https://www.arabidopsis.org/tools/go_term_enrichment.jsp. Some terms are shortened with “…”, but the full names are in Table S3. (d) Significantly down regulated processes based on all down-regulated high-fidelity protein coding genes (b), but all genes with the enriched pathways were shown. The fold of pathway enrichment, fold change and known functional interaction in the STRING database are all shown and color-coded. (e-f) Enriched biological processes based on up (e) or down (f) regulated transcription factors with medium fidelity (at least 2-fold change, p-value <0.05). GO terms were generated with PANTHER overrepresentation test by https://www.arabidopsis.org/tools/go_term_enrichment.jsp. Some terms are shortened with “…”, but the full names are in Table S4.

Out of the up- and down-regulated genes, a set of genes were obtained as high-fidelity core up- and down-regulated protein coding genes (Figure 5b, Table S3); based on these genes, 28 different pathways were up-regulated (Figure 5c and Table 3), while only 5 pathways were down-regulated (Figure 5d and Table S3). Specifically, repressed pathways include those involving very long fatty acids metabolism process, response to auxin, response to light, as well as developmental growth and cell differentiation (Figure 5d). We also examined all potential transcription factors encoded by the Arabidopsis genome and found that out of the total 1851 transcription factors, 1095 were identified, in which 88 and 29 were identified as downregulated with medium and high fidelity, respectively. Meanwhile, a total of 99 and 47 TFs were identified as up regulated with medium and high fidelity (Figure 5b, Table S4). When using all transcription factors with medium and high fidelity for pathway enrichment analysis, only a few pathways were identified (Figure 5e and f). These include upregulated pathways in response to water deprivation, salt stress, ethylene-activated signalling, as well as down-regulated pathways in trichome patterning, and epidermal cell differentiation, supporting our microscopic observation (Figures 2 and 3). It is not surprising that transcription factors in maintaining floral organ identity were identified as down-regulated because total RNAs were analyzed only from the 8-day-old seedlings.

To assess changes that occur at the protein level, we performed a proteomics analysis on 8-day-old mutant and wildtype seedlings via label-free mass spectrometry. In combination, a total of 2477 proteins were identified (Figure S5a). After normalization, 311 down- and 1183 up-regulated proteins were identified, respectively (Table S5). Of note, mass spectrometry did not detect any HSP90.7 peptide (Table S5) thus confirming the immunoblotting results (Figure 1b). Likely due to the relatively low abundance, transcription factors were not well detected by mass spectrometry analysis (Figure S5a). Pathway analysis indicated that proteins up- or down-regulated in the mutant were mainly involved in cellular metabolic pathways (Figure S5b-c, Table S5).

### Reduced auxin accumulation and impaired auxin biosynthesis pathway in *hsp90.7-1* mutant seedlings

RNA sequencing analysis indicated that only a handful pathways were significantly repressed including the one response to auxin (Figure 5d). We probed all differentially expressed genes and identified that many involved in auxin biosynthesis and conjugation were mainly upregulated, while those involved in transport and signalling were mainly downregulated (Figure 6). To confirm the role of HSP90.7 in cellular auxin homeostasis and response, we first analyzed the overall cellular auxin distribution using *DR5GUS* (Ulmasov *et al*., 1997) and *DR5rev::GFP* (Friml *et al*., 2003), the two commonly used artificial auxin sensitive reporter lines after crossing them to the *hsp90.7-1* heterozygotes.

**Figure 6.**
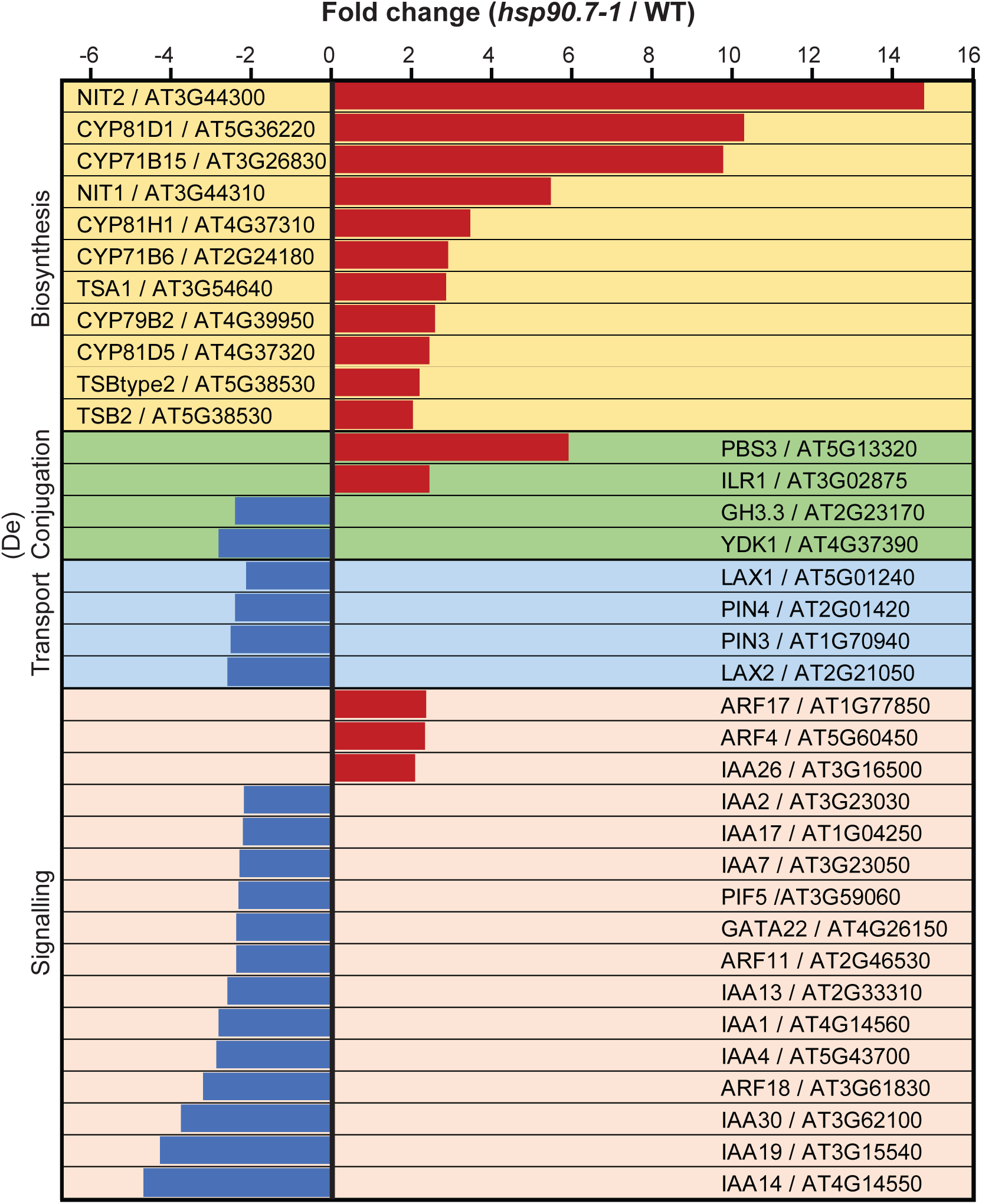
Differentially expressed protein coding genes involved in the cellular auxin homeostasis. Genes involved in auxin biosynthesis, conjugation/deconjugation, transport and response with fold change greater than 2.0 and p-value smaller than 0.05 are grouped together and shown.

In the *hsp90.7-1* mutants, *DR5GUS* had much lower expression in the SAM and root elongation zone (Figure 7a top) and *DR5revGFP* also had much less GFP signals (Figure 7b). On the other hand, *IAA2GUS*, a well characterized auxin-inducible gene (Rusak *et al*., 2010) showed a higher GUS activity in the mutant root tissues (Figure 7a bottom). We treated the corresponding reporter lines with the auxin analogue NAA (1-Naphthaleneacetic acid). It is evident that *DR5GUS* was highly inducible by NAA particularly in the SAM and the root elongation zone of the *hsp90.7-1* mutant (Figure 7a top), indicating that the auxin responsive pathway is still functioning well in the mutant. *IAA2GUS* activity in shoot and root of *hsp90.7-1* heterozygote seedlings was slightly induced by NAA, while *IAA2GUS* activity in *hsp90.7-1* homozygote mutant was strongly induced (Figure 7a bottom). This suggests hsp90.7-1 may have lower auxin content and therefore appear more sensitive to external auxin application.

**Figure 7.**
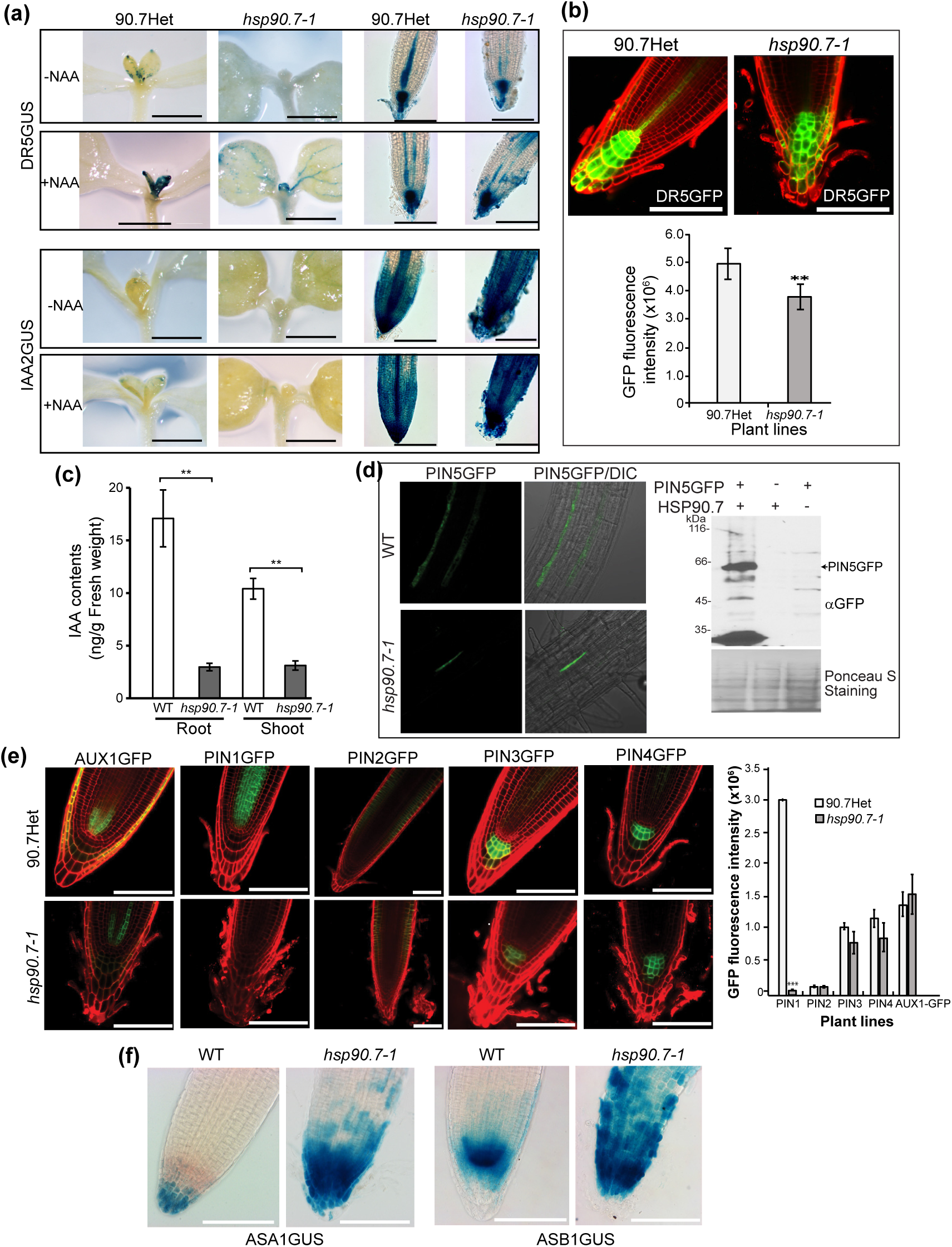
Lack of HSP90.7 leads to reduced cellular IAA content. 6- or 7-DAG seedlings were analyzed. Green is for GFP fluorescent signal and red is for propidium iodide. Scale bar: 100 μm. Quantitative analyses of the GFP signals were performed using Image J. Bars represent standard deviation. Student’s t-test were used for statistical analysis. ** indicates P < 0.005. **** indicates P < 0.0005. (a) GUS staining of root tissues from seedlings expressing DR5GUS and IAA2GUS in *hsp90.7-1* background. “+NAA” indicates seedlings were treated with 100 mg/L 1-Naphthaleneacetic acid for ∼1.5h. (b) Confocal fluorescent images of GFP signal from DR5rev::GFP construct in 90.7Het and *hsp90.7-1* mutant seedlings and the quantitative analysis (bottom). (c) Indole-3-acetic-acid (IAA) contents in root and shoot tissues of wild type and *hsp90.7-1* mutant. (d)PIN5GFP expression as revealed by confocal fluorescent images (left) and immunoblotting (right) in wild type and *hsp90.7-1* mutant seedlings. (e) Confocal fluorescent images of auxin flux and carriers AUX1GFP, PIN1GFP, PIN2GFP, PIN3GFP and PIN4GFP in 90.7Het and *hsp90.7-1* background and quantitative analysis (right). (f) GUS staining images for ASA1GUS and ASB1GUS markers in *hsp90.7-1* and 90.7Het roots.

Auxin has been known to induce another family of SMALL AUXIN UP-REGULATED RNA (*SAUR*) genes expression (Hagen and Guilfoyle, 2002, Stortenbeker and Bemer, 2019). Many SAUR proteins interact with a clade of type 2C protein phosphatase (PP2C) to inhibit its activity in dephosphorylating the plasma membrane proton pump ATPase, thus promoting proton pumping into apoplast and then cell wall elongation and growth (Spartz *et al*., 2014). Many plasma membrane-localized SAUR genes are also transcriptionally down-regulated in the mutant by more than 2-fold (Table S6), likely due to reduced cellular auxin content. To confirm all these reduced auxin responses, we measured the IAA content, and indeed found that the natural auxin IAA content was significantly reduced in both shoot and root tissues of the *hsp90.7-1* mutant seedlings (Figure 7c). Taken all together, our data demonstrated that the overall cellular auxin content was significantly lower, and auxin responsive genes were generally expressed at much reduced levels in the mutant tissues.

We then investigated whether local auxin transport is also impaired in the mutant and chose to analyze the localization and expression of a set of GFP-fused auxin transport proteins. They include auxin influx AUX1 and the polar auxin carrier proteins including PIN1, PIN2, PIN3, PIN4 and PIN5, which are all synthesized and translocated initially from the ER. Interestingly, only PIN1 and PIN5 appeared to be significantly repressed in the mutant (Figure 7d,e). PIN3 and PIN4 proteins were only slightly repressed, not as much as at the transcriptional level (Figure 6).

Moreover, the reduced auxin accumulation in mutant roots (Figure 7a-c) may result from reduced local auxin biosynthesis. To test this, we analyzed the *ASA1GUS* and *ASB1GUS* activity in primary root tissues. *ASA1* and *ASB1* encode for α and *β* subunits of anthranilate synthase, an enzyme in the upstream of tryptophan (Trp) biosynthesis (Spraggon *et al*., 2001). Interestingly, there was a much higher GUS activity from both *ASA1GUS* and *ASB1GUS* in the mutant (Figure 7f), indicating Trp biosynthesis was up-regulated, in agreement with the RNA sequencing data that indole-containing biosynthesis pathway was up-regulated (Figure 5c), as well as with the proteomics analysis showing proteins involved in tryptophan metabolism processes were also enriched in the *hsp90.7-1* mutant (Figure S5b). These data support the notation that biosynthesis of Trp, the auxin precursor molecule is induced. Considering the overall low IAA content (Figure 7a-c), Trp-dependent auxin biosynthesis therefore might be down-regulated to achieve an overall low auxin content.

Surprisingly, *NIT1* and *NIT2,* two genes in the auxiliary post-acetaldoxime Trp-dependent auxin biosynthesis pathway (Lehmann *et al*., 2017) were both up-regulated by more than 2-fold (Figure 6). This prompts that the primary TAA-YUCCA auxin biosynthesis pathway (Mashiguchi *et al*., 2011) might be significantly down-regulated. There are three characterized TAA1-family genes, *TAA1*, *TAR1*, *TAR2* (Stepanova *et al*., 2008), that encode tryptophan aminotransferase and convert Trp to indole-3-pyruvic acid (IPyA) which is further oxidized by 11 YUC family IPyA monooxygenase to IAA (Blakeslee *et al*., 2019). While *TAA1* was detected as not significantly changed, *TAR2* was down-regulated by almost 2-fold in the mutant (Table S2), thus supporting the overall observation that the primary Trp-dependent TAA-YUCCA auxin biosynthesis pathway is repressed. Likely owing to their very low expression levels, our RNA sequencing analyses did not detect any of the YUC family genes. In wild type plants, the YUC genes were differentially expressed and *YUC2*, *YUC3*, *YUC5*, *YUC6* and *YUC8* are the most widely expressed in shoot and root tissues (Figure S6).

We therefore analyzed the expression of these few widely genes by RT-qPCR. *TAA1*, *TAR2* and *PIN1* were also analyzed to validate the RNA sequencing data. In agreement with the bulk RNA sequencing analyses, *TAA1*, *TAR2*, and *PIN1* had very similar up- and down-regulation patterns in the mutant from RT-qPCR analyses (Figure 8a). As expected, although *YUC6* expression was not much changed in the mutant seedlings *YUC2*, *YUC3, YUC5* and *YUC8* were downregulated (Figure 8b), indicating that the primary TAA-YUCCA pathway was indeed repressed in the mutant. Taken all together, our analyses supported a model that knockout of ER-localized HSP90 family protein HSP90.7 shows a pleotropic effect in Arabidopsis resulting in a seedling lethality phenotype which is likely a direct result of impaired cellular auxin biosynthesis and homeostasis. In summary, relevant genes involved and especially those affected and detected in our study were schematically depicted (Figure 8c), to highlight the overall impact of HSP90.7 on the cellular auxin homeostasis.

**Figure 8.**
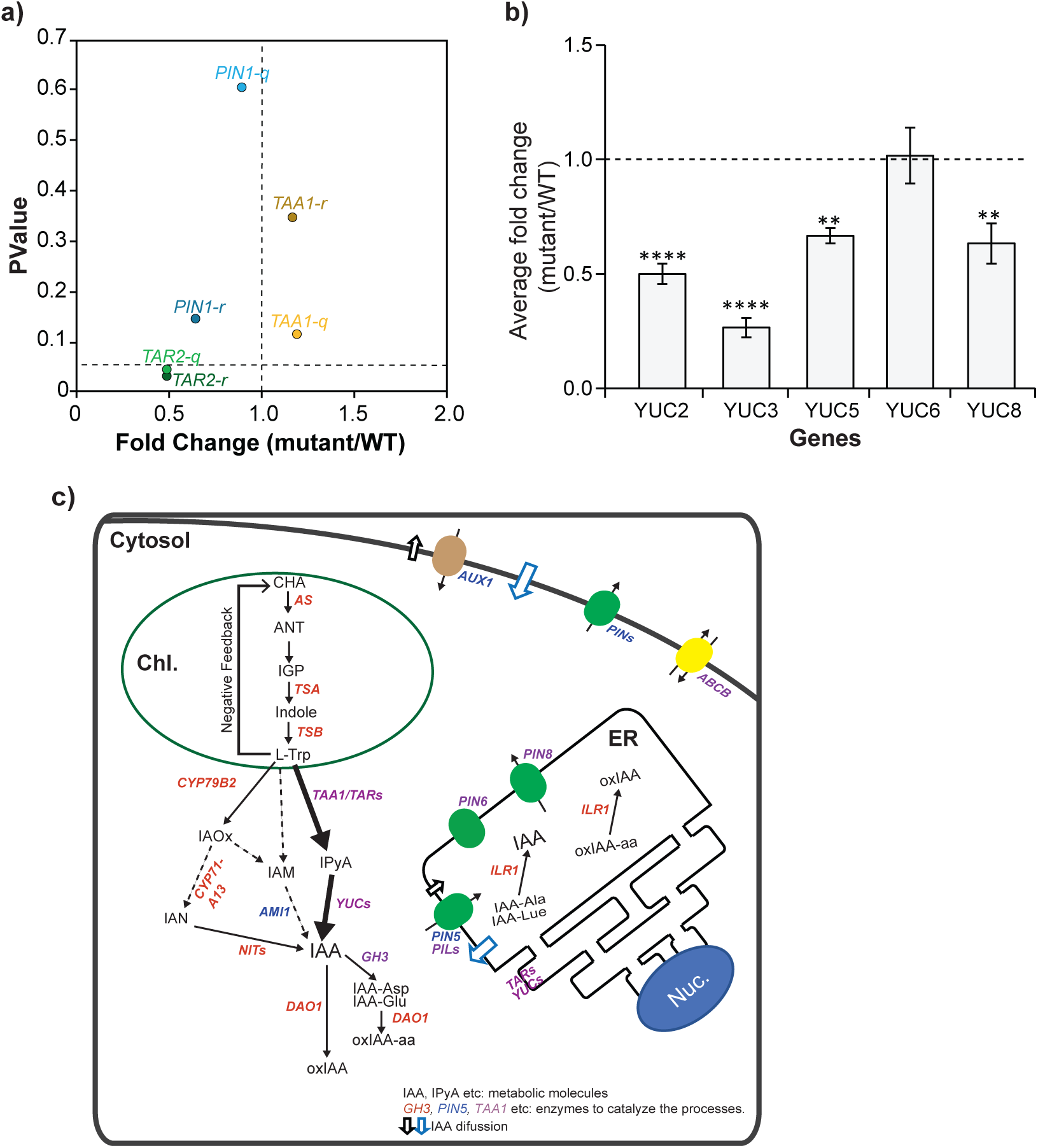
The expression of representative genes in the cellular primary auxin biosynthetic pathway. Transcript levels of TAA-YUC auxin biosynthesis genes in the *hsp90.7-1* mutant via RT-qPCR. Relative expression in the WT (control) was set to 1 for reference. Expression level of target genes was normalized to *ACT2*. (a) Average fold-change in genes that were previously detected by RNA sequencing analysis. Note: “-r” represents RNA sequencing data and “-q” represents RT-qPCR data. (b) Average fold-change of representative YUC genes Error bars represent the standard deviation from three biological replicates, each with three technical repeats. Statistical analysis was performed using the Student’s T-test. *indicates P < 0.05. **indicates P < 0.005. *** indicates P < 0.0005. (c) Schematic diagram of the Trp-dependent indole-3-acetic-acid (IAA) biosynthesis, including directional IAA transport and down-stream metabolism. Tryptophan (L-Trp) biosynthesis takes place in the chloroplast (Chl.) and the following downstream biosynthesis, conjugation and oxidation pathways mostly take place in the cytosol. Some IAA biosynthesis also takes place in the endoplasmic reticulum (ER) along with IAA-amino acid (IAA-aa) hydrolysis. Metabolic molecules are indicated in black. Enzymes are indicated in red (up-regulated), blue (down-regulated), or purple (not clear). Pathways that are not well defined are indicated by dashed arrows. [(Adapted from (Perez-Llorca *et al*., 2019)] Abbreviations: ABCBs, ATP-binding cassette transporter; ANT, Anthranilate; AUX1, AUXIN1/LIKE-AUX1 (AUX1/LAX); DAO1, DIOXYGENASE FOR AUXIN OXIDATION 1; GH3, GRETCHEN HAGEN3; IAA, indole-3-acetic acid; IAM, indole-3-acetamide; IAN, indole-3-acetonitrile; IAOx, indole-3-acetaldoxime; IGP, indole-3-glycerol phosphate; ILR1, IAA-LEUCINE RESISTANT1; IPyA, indole-3-pyruvic acid; NITs, Nitrilases; oxIAA, 2-oxindole-3-acetic acid; oxIAA-aa, 2-oxindole-3-acetic acid-amino acid; PILS, PIN-likes; PIN, PIN-FORMED; TAA1, TRYPTOPHAN AMINOTRANSFERASE OF ARABIDOPSIS 1; TARs, TAA1-RELATED proteins, tryptophan aminotransferases; TSA/TSB, tryptophan synthase subunit A/B; YUCs, flavin-containing monooxygenases.

## DISCUSSION

ER-localized HSP90 family chaperones have been identified and recognized particularly for higher multicellular organisms and absent in lower unicellular organisms like baking yeast. It has been postulated that HSP90 paralogs within the ER are required for the biogenesis of certain receptor proteins, such as TLR, immunoglobulin which are only available for multicellular organisms (Kim *et al*., 2021). Plant ER-located HSP90.7 is required for the maintenance of the SAM by regulating the WUSCHEL-CLAVATA pathway involving receptor kinase CLV1 and the ligand CLV3 (Ishiguro *et al*., 2002). In this study, we identified and investigated for the first time a complete HSP90.7 knockout mutant in Arabidopsis and confirmed *CLV1* was down regulated in the mutant. We further provided evidence that HSP90.7 is also required for RAM niche maintenance, male and female fertility, and for cellular endomembrane system homeostasis. Particularly, we confirmed HSP90.7 is required for cellular PIN1 and PIN5 protein accumulation. Though PIN1 and PIN5 are not receptor like kinases, they export auxin and help establish appropriate hormonal cues to mediate cell to cell communication (Krecek *et al*., 2009). We also performed high throughput analysis to identify differentially expressed genes in the *hsp90.7-1* mutant at both the RNA and protein levels and provided a comprehensive array of HSP90.7 functional interactors. Our study therefore established that the ER-localized HSP90 family protein plays a globally essential role in higher flowering plants.

In addition to its previously reported role in apical meristem niche maintenance, HSP90.7 is also required for the development of trichomes (Figure 4b), which typically develop in and act as a hallmark of vegetative leaves (Pesch and Hulskamp, 2009). The initiation and maturation of trichomes requires sequential structural changes of epidermal cells and is controlled by a set of well conserved transcription factors like MYB23 and the trichome initiation complex GL1-GL3-TTG1 in both unicellular and multicellular trichomes (Han *et al*., 2022, Wang *et al*., 2022). Particularly, expression of GLABRA2 (GL2) which is controlled by the initiation complex promotes trichome development and repression of GL2 inhibits trichome development. In the *hsp90.7-1* mutant, MYB23, TTG2, and particularly GL2 were all significantly inhibited (Table S2) resulting in repressed trichome differentiation. Interestingly, it has been well documented that trichome development and root hair development share some core transcription factors, but GL2 plays a completely opposite role, such that expression of GL2 inhibits root hair development while repression of GL2 promotes root hair development (Hulskamp, 2004). The completely opposite phenotypes in trichome and root hair development of *hsp90.7-1* mutant (Figure 4 and Figure S3c) agree well with the RNA sequencing data.

HSP90.7 is an ER-localized molecular chaperone, and it is generally thought that comprise of the protein function would elicit unfolded protein response in ER. In *hsp90.7-1* mutant, we observed up-regulation of BiP protein (Figure 3f), a hallmark of the ER stress (Gardner *et al*., 2013), suggesting the mutant seedlings experience ER-associated unfolded protein response (UPR), which normally induces ER-associated protein degradation (ERAD) or autophagy (ER-Phagy) (Li and Sun, 2021, Yang *et al*., 2021). Nevertheless, we noted although some other ER-located chaperones like CNX, and CRT are also up-regulated at both transcriptional and translational levels (Table S2 and S5), genes associated with ERAD in the mutant like *OS9, HRD1, HRD3A, RIN2, RIN3, SUD1*, *SEL1L* are not differentially expressed (Table S2), suggesting that knockout of HSP90.7 may not induce the ERAD pathway. Much accumulated ER-lumen marker protein (Figure 3E) indicated that the cellular endomembrane system is significantly altered and whether the increased BiP protein in the mutant is mainly a result of increased cellular ER network remains further investigation. Additionally, GRP94 depletion in mammalian cells triggers UPR, but does not induce expression of protein disulfide isomerase (PDI) proteins (Mao *et al*., 2010), while HSP90.7 knockout results in much induced PDI-like proteins (PDIL1-1, PDIL1-2, PDIL1-4, PDIL2-2, PDIL2-3, PDIL1-3) (Table S5). This suggests different roles of the ER-located HSP90 paralogs in plant and mammalian cells.

We identified the auxin responsive pathway as one of the only few that are repressed in the *hsp90.7-1* mutant (Figure 5d). We measured a significantly reduced IAA content in both root and root tissue (Figure 7c) as well as reduced response from auxin response marker proteins (Figure 7a,b). Cellular auxin homeostasis involves auxin biosynthesis, catabolism/conjugation, long and short distance transport, and protonated neutral auxin is also readily diffusible across membranes. Although it is unlikely to attribute a single or a major process to the reduced IAA level in the mutant seedlings, we provided evidence that, in the mutant, the major TAA-YUCCA auxin biosynthesis pathway was reduced, and that PIN1 and PIN5 expressions were also significantly reduced. PIN1 is a primary protein for polar auxin transport and compromise of cytosolic HSP90 isoforms also affects PIN1 polar distribution (Samakovli *et al*., 2021). However, PIN1 function might be more impacted by the ER-localized HSP90.7 as they do have the chance to physically interact within the endomembrane system. Whether PIN1 and PIN5 directly interact with HSP90.7 during their biogenesis and act as HSP90.7 clients remains mysterious.

Additionally, multiple PIN family members like PIN1, PIN2 and PIN3 are localized on the same plasma membrane and they form homo- or heterodimers in vivo, and their stability and polar auxin transport activity can be further regulated by endogenous flavonols (Teale *et al*., 2021), phosphorylation (Huang *et al*., 2010) and inhibited by 1-naphthylphthalamic acid (NPA) which binds to the same pocket as natural auxin IAA but with much higher affinity (Abas *et al*., 2021, Su *et al*., 2022, Yang *et al*., 2022). However, not much is known on how molecular chaperones are associated with these proteins during their folding and targeting processes to the dedicated membranes. HSP90.7 might be one needed for in vivo biogenesis of these PIN proteins. Lastly, we did not examine further the function of representative auxin catabolism related genes, but, the *DIOXYGENASE FOR AUXIN OXIDATION 1* (DAO1), and *IAA-LEUCINE RESISTANT 1* (ILR1) which hydrolyzes oxIAA-amino acids into oxIAA (Hayashi *et al*., 2021) were all observed to be upregulated (Table S2), suggesting the GH3-DAO-ILR1 auxin conjugation pathway is induced and may partly attribute to the reduced auxin content in the *hsp90.7-1* mutant. In summary, our study provided many potential targets to understand how those proteins regulating cellular auxin homeostasis themselves can be regulated at the posttranslational levels by molecular chaperones.

## EXPERIMENTAL PROCEDURES

### Plant material and growth conditions

*Arabidopsis thaliana* (Col-0) was used in this study as the wildtype plant and seeds were obtained from the Arabidopsis Biological Resource Center, ABRC (Columbus, OH). All wildtype and transgenic plant lines used this study are summarized in Table S7. For in vitro culture, seeds were surface-sterilized and sown on half strength Murashige and Skoog (MS) media containing 1% (w/v) sucrose and 0.8% agar, without or with 10 µg mL^−1^ BASTA, 20 µg mL^−1^ kanamycin, or 25 µg mL^−1^ hygromycin for selection. Seeds were stratified in the dark at 4°C for 3 days and then incubated at 22°C, 16h light/8h dark photoperiod with illumination of 110 µmol.m^−2^.s^−1^. For root growth assay, seeds were laid out on petri dishes vertically to promote root growth on the surface of the agar. For late stage growth and development, seedlings were transferred to soil and grown in Conviron ATC60 growth chamber at 22°C, 16 h light/8 h dark photoperiod with illumination of 110 µmol.m^−2^.s^−1^.

### Plasmid constructions and plant transformation

To make a construct for HSP90.7 expression in plant under its endogenous promoter, the CaMV 35S promoter in pGWB502Ω (Nakagawa *et al*., 2007) was substituted with the 1022 bp predicted HSP90.7 promoter sequence together with the 5’UTR region (https://agris-knowledgebase.org/AtcisDB/). To this end, the promoter fragment was amplified and cloned between HindIII and XbaI sites using In-Fusion® HD Cloning Kit (Takara), generating pGWB502-HSP90.7pro. A previously generated Gateway entry vector encoding a C-terminally FLAG-tagged HSP90.7 coding region (Chong *et al*., 2015) was introduced to generate pGWB502omega-HSP90.7pro-HSP90.7FLAG by LR reaction. A 1022bp HSP90.7 promoter fragment completed with the 5’UTR and 21bp downstream of ATG was cloned into pBI101.1 between HindIII and BamHI to generate a GUS reporter construct. To generate an mCherry tagged HSP90.7, an NcoI site was introduced into the FLAG-tagged HSP90.7 entry vector before KDEL by site-directed mutagenesis, and the mCherry encoding sequence was amplified from ER-RK (Nelson *et al*., 2007) and then inserted into the NcoI site. All primers used to make different constructs, PCR genotyping and RT-qPCR are included in Table S8. All binary vectors were transformed into *A. tumefaciens* GV3101 and then used to transform *Arabidopsis thaliana* Col-0 by floral dip method (Clough and Bent, 1998).

### GUS histochemical staining

Young seedlings or pollens were incubated in GUS staining buffer (1 mM EDTA pH 8.0, 5 mM potassium ferricyanide, 5 mM ferrocyanide, 100 mM sodium phosphate pH 7, 0.1% Triton X-100) containing 1 mg.mL^−1^ X-Gluc (5-bromo-4-chloro-3-indolyl-beta-D-glucuronic acid) at 37 °C, and then depigmented with 95% ethanol and stored in 50% glycerol. The stained tissues or cells were analyzed with Zeiss AxioScope epifluorescence or light dissecting microscopes.

### Root cortex cell number count and cell size measurement

The number of outer cortical cells within the root apical meristem (RAM) were counted from the quiescent center (QC) until the onset of rapid elongating cells (REZ) using a criteria as previously described (Cajero Sanchez *et al*., 2018). The number of outer cells in the rapid elongation zone were counted from the onset of rapid elongation until the outer cortical cell closest to the epidermal cell with the first root hair bulge. Using the same criteria, the average length of the cortical cells within the RAM and REZ were measured using ZEN 3.0 SR in μm.

### In vitro pollen germination

Freshly opened anther-dehisced flowers were collected for *in vitro* pollen germination as previously described (Fan *et al*., 2001) with some modifications. Flowers were dipped on a drop of growth medium [5 mM MES (pH 5.8 adjusted with TRIS base), 1 mM KCl, 0.8 mM MgSO_4_, 1.5 mM boric acid, 10 mM CaCl_2_, 5% (w/v) sucrose, 15% (w/v) PEG4000] on the surface of a glass slide. The pollen grains were germinated at 28 °C with 2 to 24 h of incubation as needed.

### SDS-PAGE and immunoblotting analysis

Plant tissues were grinded in liquid nitrogen and boiled in 2x SDS-sample buffer. Protein concertation was normalized to the fresh weight (w/v) or estimated using Bradford protein assay reagent (BioRad). Protein samples were subjected to SDS-PAGE followed by a transfer onto nitrocellulose membrane (PALL BioTrace™; 66,485) and immunoblotted with specific antibodies, including anti-HSP90.7 polyclonal antibody (Chong *et al*., 2015), anti-FLAG monoclonal antibody (F3165; Sigma), and anti-GFP polyclonal antibody (SAB4301138; Sigma), anti-BiP antibody (E1811; Sant Cruz).

### Light and electron microscopy and imaging

Confocal images were obtained using a Zeiss 510 laser scanning microscope (LSM). When needed, root tissues were stained in 0.01 μg/mL propidium iodide in ½ MS liquid media for 5-7 minutes before visualization. For GFP signal quantification, samples were visualized on the same day and under identical confocal settings, and fluorescence signals were quantitated with ImageJ software (http://rsbweb.nih.gov/ij/). To visualize high resolution cell structures, 6- to 8-day-old root or shoot tissues were fixed in 2% (w/v) glutaraldehyde in 0.1 M Sorensen’s phosphate buffer (pH 7.2) and 2% (w/v) sucrose at room temperature for 2 hours, then dehydrated in an ethanol series before getting embedded in Spurr resin. One-micron semi-thin cross sections were cut with glass knives and stained in 1% Toluidine Blue and 1% borate acid. Sections were imaged using Zeiss AxioScope epifluorescence microscope. For transmission electron microscopy, samples were then postfixed with 2% OsO4 for 6 hours and dehydrated in an ethanol series before getting embedded in Spurr resin. Ultrathin sections (50 nm thick) were cut with glass knives and stained with uranyl acetate and lead citrate before observation with Hitachi H7500 electron microscope. For scanning electron microscopy, post fixation was in 1% (w/v) OsO4 for 1 hour at room temperature, dehydrated in an ethanol series and critical point dried with CO_2_ (Leica EM CPD300), then sputter-coated with osmium before observation with Hitachi S530 scanning electron microscope.

### RNA sequencing and GO term enrichment analyses

Total cellular RNA was extracted from 8-day-old wildtype or mutant seedlings grown on 1/2MS medium with RNeasy Mini Kit and QIAshredder (Qiagen). Three independently prepared samples with RNA integrity number greater than 8.0 from each line were sequenced and analyzed at The Centre for Applied Genomics at SickKids Hospital (Toronto, Canada). The trimmed reads were screened using FastQ-Screen v.0.10.0. The variance stabilized counts were used in principal component analysis (PCA) to check the sample quality and expressed genes were searched against TAIR10 database. Raw gene counts were loaded and sample-normalized using DESeq v.1.18.0. Two-condition differential expression was done with the edgeR R package v.3.22.3 (Robinson *et al*., 2010). The data set was filtered to retain only genes whose FPKM > 2 in at least 2 samples, thus removing genes that are not expressed, or expressed at a very low level. Differentially expressed genes with at least 2-fold changes were used for GO term enrichment analysis with PANTHER™ knowledgebase (Thomas *et al*., 2022).

### Label-free mass spectrometry

The total soluble proteins from fresh 8-day-old seedlings were prepared and analyzed by liquid chromatography coupled with tandem mass spectrometry (LC-MS/MS) as previously described (Bonea *et al*., 2021) at the SPARC BioCentre at SickKids Hospital (Toronto, Canada). Scaffold 4.8.8 (Proteome Software Inc.) was used to refine peptide and protein identifications. Proteins with over 99.0% probability and at least two exclusive unique peptides from at least one sample were accepted for further examination. Differentially accumulated proteins in the mutant were analyzed for enriched pathways with go:profile based on KEGG pathway database (Raudvere *et al*., 2019).

### RT-qPCR analysis

Real-time quantitative PCR (RT-qPCR) was performed as previously described using the same RNA purified for RNA sequencing analysis. Superscript first-strand synthesis system (Invitrogen) was used for the synthesis of cDNA from 1 mg of total RNA. cDNA (10ng) was added to iQSYBRE Green qPCR supermix (Bio-Rad) with appropriate primers (Table S2). Data analysis was performed with Thermofisher QuantStudio3 thermocycler and Design and Analysis software (version 2.6).

### IAA content measurement

Procedures for extraction, purification, and measurement of IAA by a liquid chromatography-electronspray ionization-tandem mass spectrometry (LC-ESI-MS/MS; Agilent 6410 TripleQuad LC/MS system) are previously described (Lu *et al*., 2015). D2-IAA (Olchemim, CZ) was added as an internal standard before purification.

### Statistics analysis

Two-tailed, unpaired Student’s t-test assuming unequal variance were used to check for significance (p < 0.05) in the comparison of control and treatment group. For experiments having two or more control groups, two-way analysis of variance (ANOVA) assuming interaction between factors (i.e. genotype and chemical treatment) and Tukey’s honesty significant difference (HSD) tests were used to check for significance (p < 0.05) in multiple comparisons of means.

## ACCESSION NUMBERS

Genes described in this work are *HSP90.7/AT4G24190; BIP1/AT5G28540; PIN1/AT1G73590; PIN5/AT5G16530; TAA1/AT1G70560; TAR2/AT4G24670; YUC2/AT4G13260; YUC3/AT1G04610; YUC5/AT5G43890; YUC6/AT5G25620; YUC8/AT4G28720; CLV1/AT1G75820; CLV2/AT1G17240; CLV3/AT2G27250; WUS/AT2G17950*

## Abbreviations

HSP90.7: Arabidopsis ER-localized heat shock protein 90
RNAseq: Bulk RNA sequencing analysis
IAA: indole-3-acetic acid
GFP: green fluorescent protein
PI: propidium iodide
GUS: beta-glucuronidase
TEM: transmission electron microscopy
SEM: scanning electron microscopy
SAM: shoot apical meristem
RAM: root apical meristem

## ACKNOWLEDGMENTS

We would like to acknowledge Durga Acharya and Bruno Chue from the Centre for the Neurobiology of Stress (CNS) at the University of Toronto Scarborough for their assistant in performing light and electron microscopic analysis, to acknowledge Ms. Ayako Nambara for her technical assistance in IAA analysis. The authors would also like to thank Dr. Jiri Friml (Institute of Science and Technology, Austria) and Dr. Xuejun Hua (Zhejiang Industrials University, China) for providing a variety of the GFP and/or GUS reporter lines. This research was funded by a NSERC Discovery Grant to R.Z. (RGPIN-2019-07060). B.M. and J.N. were also supported by the University of Toronto graduate scholarships.

## AUTHOR CONTRIBUTIONS

JN performed most of the experiments including root and shoot apical microscopic analysis, major auxin related marker lines analyses and qPCR, and wrote the draft manuscript. BM cloned the endogenous HSP90.7 promoter and made constructs to complement the mutant alleles. HH analyzed heterozygote fertility and performed in vitro pollen germination assays. MM analyzed GUS fusion protein expressions. DB screened and confirmed the *hsp90.7-1* mutant allele and some quantitative RNA expression analysis. EN performed IAA content measurement. RZ conceptualized the project, supervised, requested funding, validated data, and wrote the manuscript. All authors reviewed and edited the manuscripts.

## SUPPORTING INFORMATION

Figure S1. PCR genotyping of hsp90.7-1 mutant allele.

Figure S2. HSP90.7 tissue specific expression from high throughput analysis.

Figure S3. *hsp90.7-1* mutant has defective primary root growth and root apical meristem organization.

Figure S4. HSP90.7 transcripts from RNA sequencing analyses.

Figure S5. Summary of differentially accumulated proteins and associated pathways in *hsp90.7-1* mutant.

Figure S6. Heatmap of YUC family gene expression in Arabidopsis tissues.

Table S1. Segregation ratio of HSP90.7 mutants.

Table S2. DEGS from RNAseq analysis.

Table S3. Up- and down-regulated core high fidelity gene sets from RNAseq analysis.

Table S4. Up- and down-regulated transcription factors.

Table S5. Differentially accumulated proteins identified by label-free LC-MS/MS.

Table S6. Auxin responsive genes identified by bulk RNA sequencing.

Table S7. Plant lines used in this study.

Table S8. Primers used in this study.

