## Supplemental Figure 1-6 for "Knockout of endoplasmic reticulum localized molecular chaperone HSP90.7 impairs seeding development and cellular auxin homeostasis in Arabidopsis"

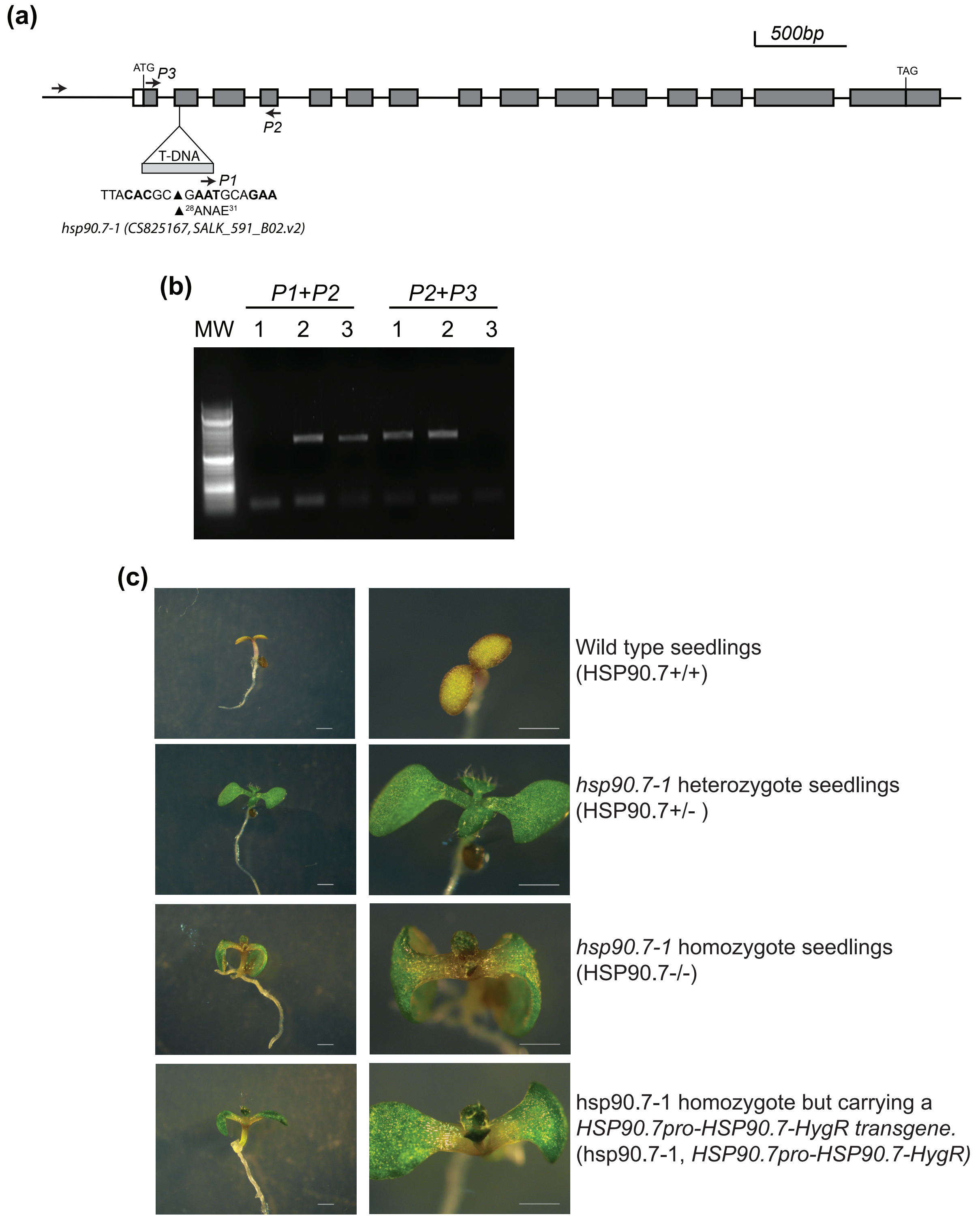


**Figure S1. PCR genotyping of *hsp90.7-1* mutant allele.**
**(a)** Same as Figure 1a. Primers P1, P2 and P3 are pCSA110-LB3, HSP90.7E138R and HSP90.7L16F, respectively, and also included in Supplemental Table 8.
**(b)** PCR genotype typing results for wild type (1), the heterozygote (2) and homozygote (3) of the hsp90.7-1 mutants using primers P1, P2 and P3 as indicated in (a).
**(c)** Representative seedlings grown on MS media with herbicide Basta® (Glufosinate) for 6 days.

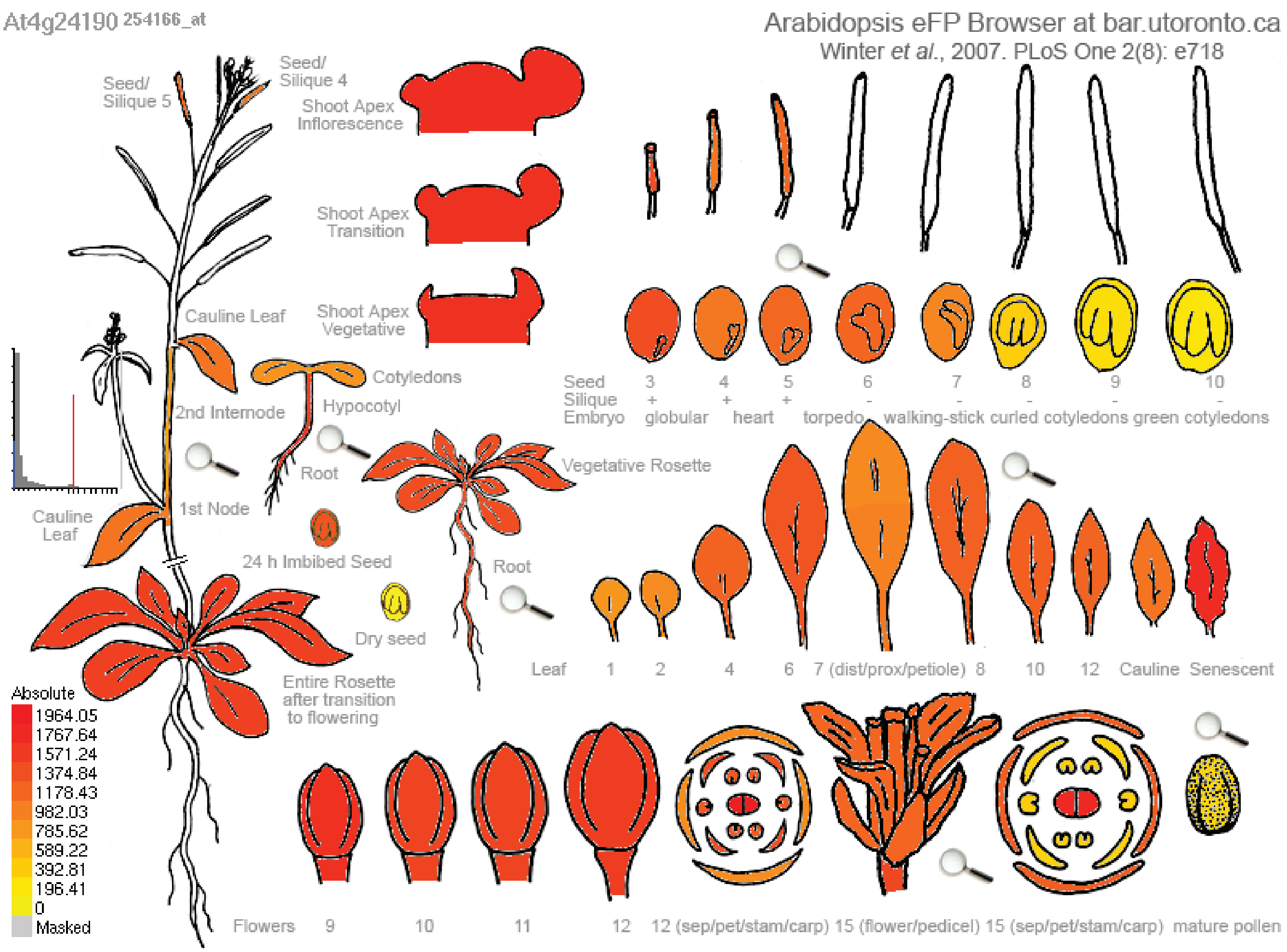


**Figure S2. HSP90.7 tissue specific expression from high throughput analysis.** Tissue specific expression of HSP90.7 as revealed by BAR (http://bar.utoronto.ca/). HSP90.7 is widely expressed in most tissues and highly expressed in apical meristems.

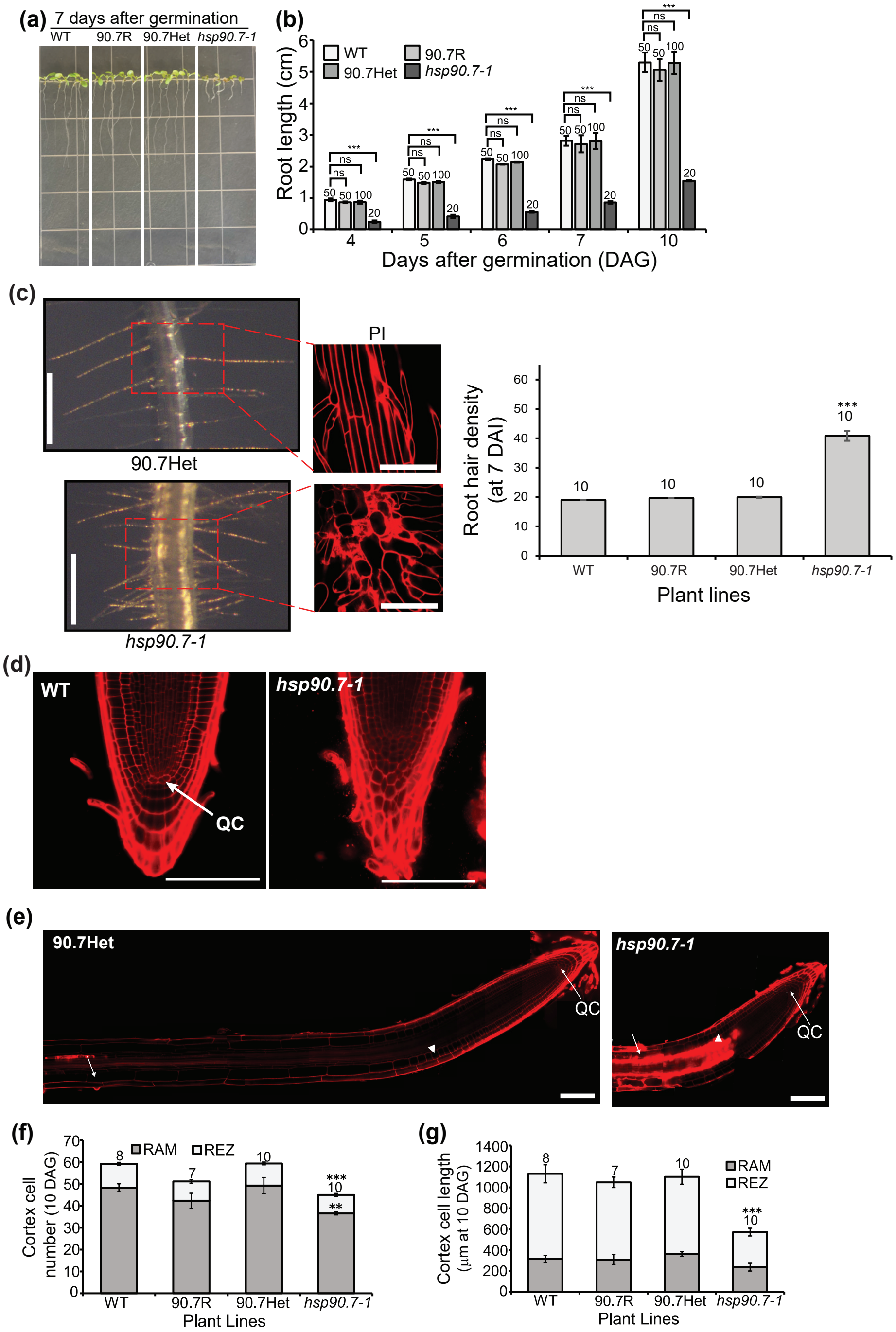


**Figure S3. *hsp90.7-1* mutant has defective primary root growth and root apical meristem organization.** **(a)** Seedlings grown vertically on petri dishes at 7-DAG. **(b)** Primary root lengths from different plant lines vertically grown up to 10-DAG. **(c)** Root hair density of the *hsp90.7-1* mutant seedlings. Microscopic root images for *hsp90.7-1* heterozygote (90.7Het) and homozygote (*hsp90.7-1*). The images on the top right are stained with propidium iodide. The bottom graph shows the root hair density for wild type (WT), the heterozygote, the homozygote and the homozygote rescued by an mCherry tagged HSP90.7 under the HSP90.7 endogenous promoter (90.7R). The number of roots analyzed were indicated on the top of the bars. **(d)** Root tip confocal images after staining with propidium iodide from WT and *hsp90.7-1* seedlings grown at 10-DAG. Scale bar: 100μ m. **(e)** Confocal images of root meristem zoon, rapid elongation zoon and maturation zone. 10-DAG seedlings roots were stained with propidium iodide. The beginning of rapid elongation zone and maturation zone were marked by a triangle and arrow, respectively. Quiescent center (QC) is also indicated by an arrow. Scale bar: 100 μm. **(f-g)**. Cortex cell number (e) and length (f) of primary roots of seedlings grown at 10-DAG. Error bars represent standard deviation and Student's t-test was used for statistical analysis. Sample numbers for statistical analysis were indicated on top of the bars. **P < 0.005, and ****P < 0.0005.

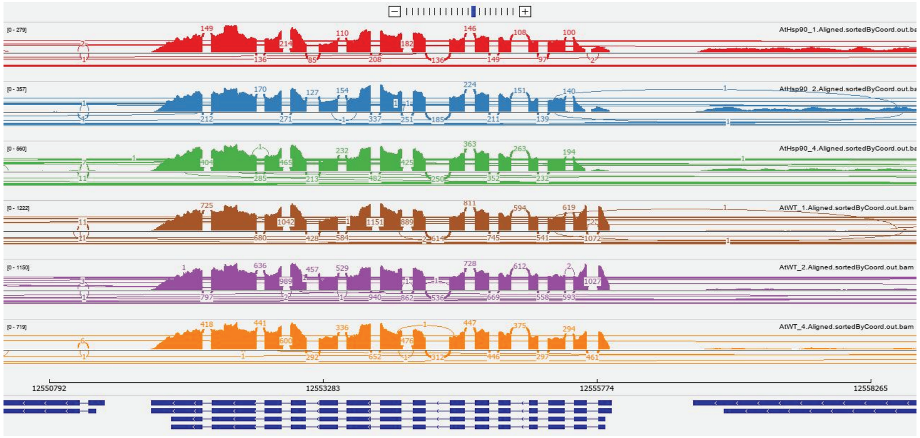

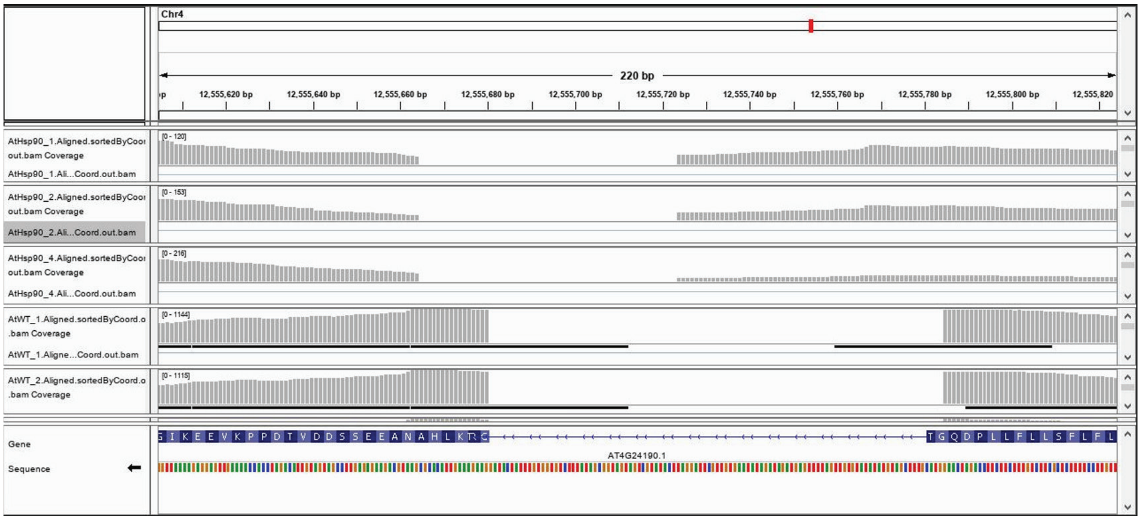

**Figure S4. HSP90.7 transcripts from RNA sequencing analyses.**
HSP90.7 transcripts in wild type and *hsp90.7-1* mutant seedlings as read by RNA sequence. The diagrams were visualized by Integrative Genome Viewer for the length of the entire gene (top) and the region focusing on the first and second exon (bottom).

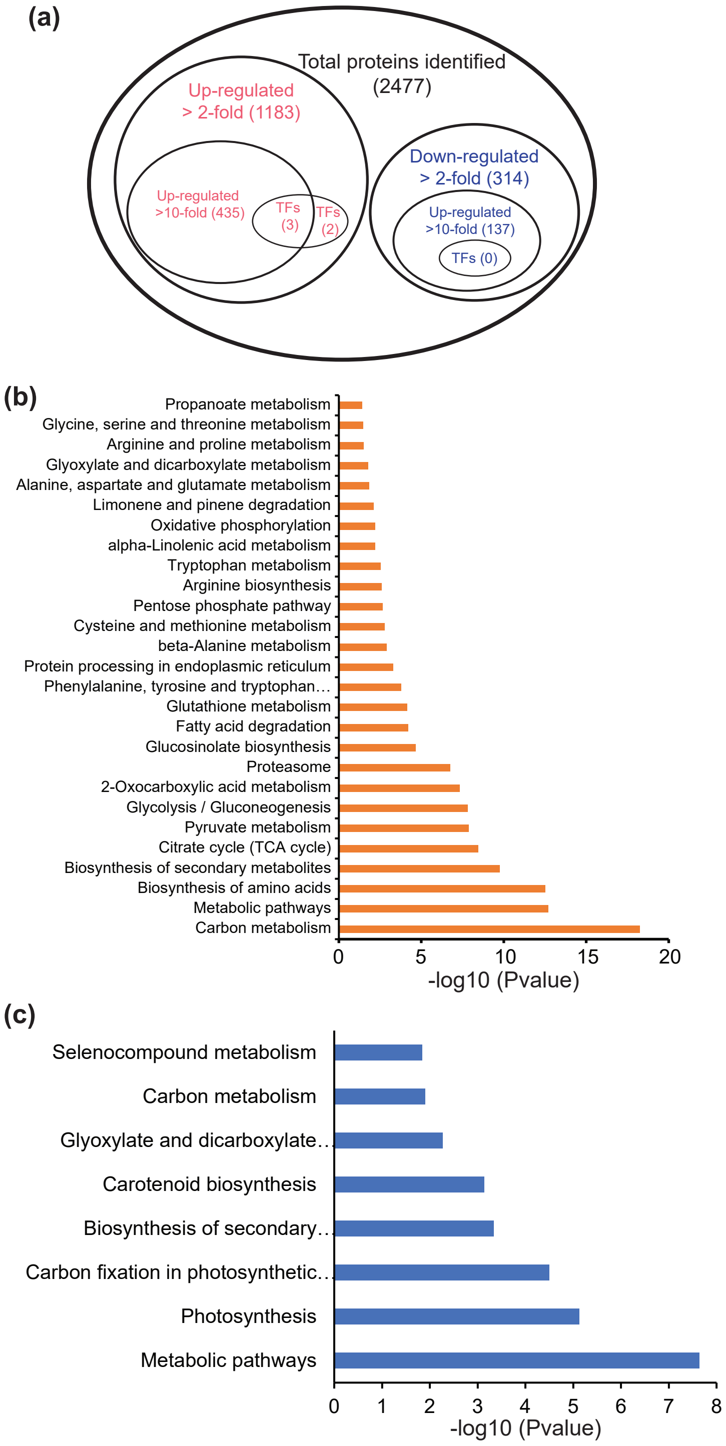


**Figure S5. Summary of differentially accumulated proteins and associated pathways in *hsp90.7-1* mutant.**

**(a)** Summary of identified proteins by label-free spectrometry of total soluble proteins in 8-DAG wild type and mutant seedlings. The raw data are indicated in supplemental Supplemental Table S5.

**(b-c)** Enriched up- and down-regulated pathways, respectively, based on proteins that are altered at least 2-fold in the mutant. GO terms based on KEGG database were analyzed by go:Profile (https://biit.cs.ut.ee/gprofiler/gost). Some terms are shortened with "…", but the full names are in Supplemental Table S5.

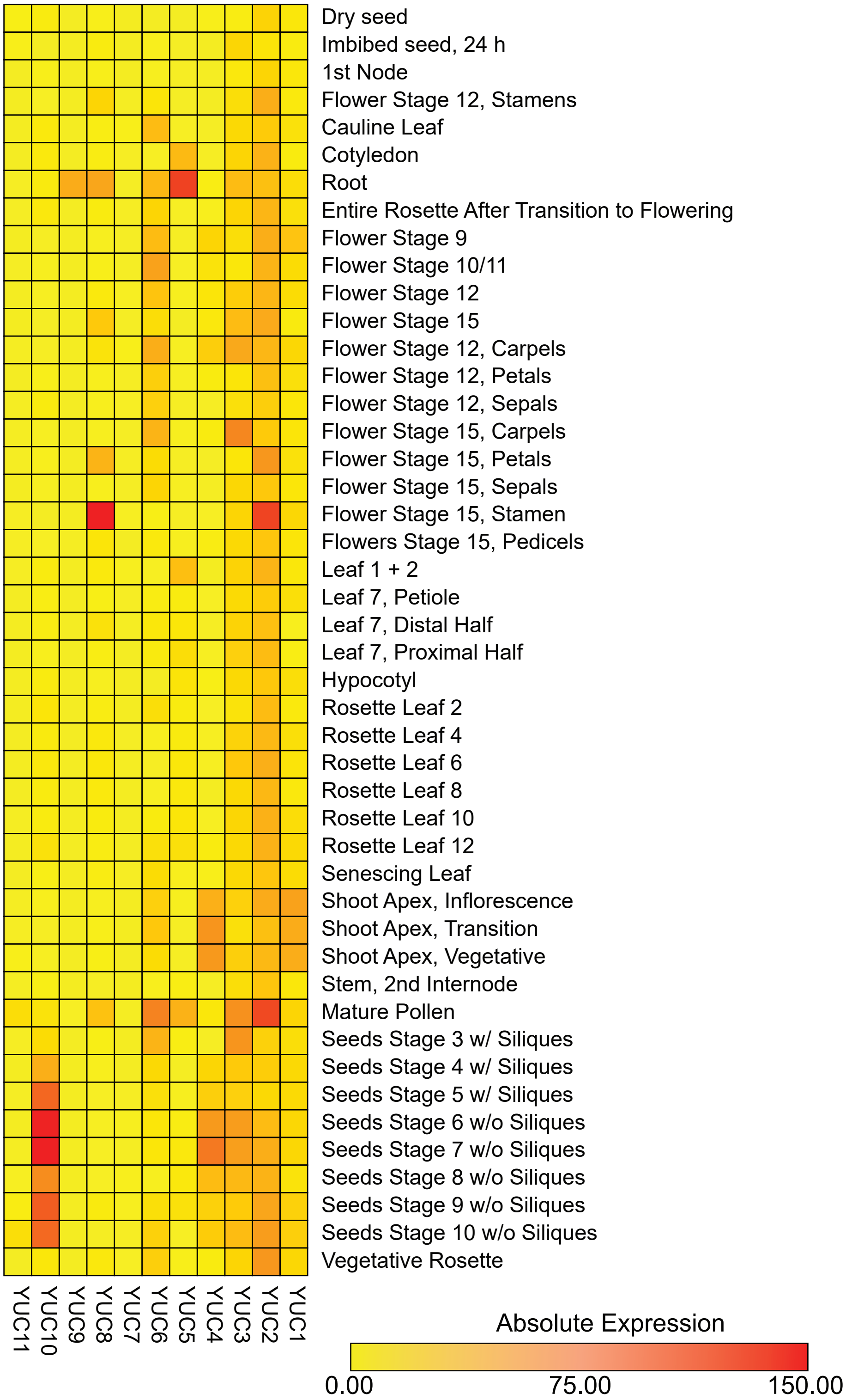


**Figure S6. Heatmap of YUC family gene expression in Arabidopsis tissues.** Tissue specific expression of 11 YUC family genes as retrieved from BAR (http://bar.utoronto.ca/) and visualized with Morpheus (https://software.broadinstitute.org/morpheus).
